# A simulation study of the ecological speciation conditions in the Galician marine snail *Littorina saxatilis*

**DOI:** 10.1101/2022.02.08.479545

**Authors:** M. Fernández-Meirama, E. Rolán-Alvarez, A. Carvajal-Rodríguez

## Abstract

In the last years the interest on evolutionary divergence at small spatial scales has increased and so did the study of speciation caused by ecologically-based divergent natural selection. The evolutionary interplay between gene flow and local adaptation can lead to low-dispersal locally adapted specialists. When this occurs the evolutionary interplay between gene flow and local adaptation could eventually lead to speciation.

The *L. saxatilis* system consists of two ecotypes displaying a microhabitat-associated intraspecific dimorphism along the wave-exposed rocky shores of Galicia. In spite of being a well-known system, the dynamics of the ecotype formation remains unclear and cannot be studied from empirical evidence alone. In this study, individual-based simulations were used to incorporate relevant ecological, spatial and genetic information, to check different evolutionary scenarios that could evolve non-random mating preferences and finally may facilitate speciation.

As main results, we observed the evolution of intermediate values of choice which matches estimates from empirical data of *L. saxatilis* in Galician shores and coincides with previous theoretical outcomes. Also, the use of the mating correlation as a proxy for assortative mating led to spuriously inferring greater reproductive isolation in the middle habitat than in the others, which does not happen when directly considering the choice values from the simulations. We also corroborate the well-known fact that the occurrence of speciation is influenced by the strength of selection. Taken together, this means, also according to other *L. saxatilis* systems, that speciation is not an immediate consequence of local divergent selection and mating preferences, but a fine tuning among several factors including the ecological conditions in the shore levels, the selection strength, the mate choice stringency and cost to choosiness. The *L. saxatilis* system could correspond to a case of incomplete reproductive isolation, where choice intensity is intermediate and local adaptation within the habitat is strong. These results support previous interpretations of the *L. saxatilis* model system and indicate that further empirical studies would be interesting to test whether the mate choice mechanism functions as a similarity-like mechanism as has been shown in other littorinids.

## Introduction

Speciation caused by ecologically based divergent natural selection can occur at small spatial scales. Microgeographic adaptation occurs when neighbour groups of individuals have adaptively diverged (Richardson et al., 2014) and this adaptation at small spatial scale may happen despite a high potential for mixing within the dispersal area. This idea has produced controversy between theoretical studies that show that natural selection can overcome the effects of gene flow (Dieckmann and Doebeli, 1999; Gavrilets, 2004; Barluenga et al., 2006; Hey, 2006; Bolnick and Fitzpatrick, 2007; Rolán-Alvarez, 2007; Debarre and Gandon, 2011; Weissing et al., 2011; Savolainen et al., 2013; Butlin et al., 2014; Richardson et al., 2014; Getz et al., 2016; Rettelbach et al., 2016; Servedio and Boughman, 2017; Kopp et al., 2018) and those that admit that this is feasible but that the conditions necessary for it to occur make it difficult to observe in the real world. (Jiggins, 2006; Babik et al., 2009; Meyers et al., 2012; Martin, 2013; Savolainen et al., 2013; Martin et al., 2015; Fernández-Meirama et al., 2017a; Foote, 2018).

There are various plausible scenarios for the evolutionary interplay between gene flow and local adaptation. They may give rise to a monomorphic population, or to generalists adapted to different habitats, or to polymorphic subpopulations of locally adapted specialists, which, in case of low gene flow, could lead to speciation (Coyne and Orr, 2004).

However, for local adaptation to occur, a gene flow-reducing mechanism, such as non-random mating due to mate choice, may be necessary in addition to selection. Furthermore, to detect mate choice, it is important to take into account the true spatial scale at which mate choice occurs to avoid a version of the so-called Simpson’s paradox in which the mating pattern existing in some data sets will disappear or be reversed when the groups are combined. Thus measuring mate choice at the correct scale is key to understanding this evolutionary process (Rolán-Alvarez et al., 2015b; Estévez et al., 2018).

Local adaptation and speciation through non-random mating are expected to occur when the traits involved in divergent selection and mate choice are the same, the so-called magic trait (Gavrilets, 2004; Thibert-Plante and Gavrilets, 2013; Servedio and Boughman, 2017; Richards et al., 2019).

Because of the long time scale necessary for local adaptation and mate choice evolution under gene flow, simulation has become an essential part of speciation research. Recent efforts on the study of the evolution of reproductive isolation focused on the complex interactions that emerge in the presence of local adaptation, mate choice and sexual selection under different genetic structure, migration model, etc. (Sadedin et al., 2009; Thibert-Plante and Hendry, 2009, 2011; Cotto and Servedio, 2017; Sachdeva and Barton, 2017; Perini et al., 2020).

The present work is based on *L. saxatilis* as model organism and used individual-based simulation to identify some relevant parameters, as local selection intensity, hybrid zone environment and demography, that can influence the evolution of local adaptation and non-random mating preferences. The *L. saxatilis* model case relies on the existence of two main ecotypes that have evolved by natural selection to adapt to distinct microhabitats in different geographical regions in parallel (see next section). The two ecotypes have evolved different average sizes, morphologies, behaviour and physiologies, and may potentially show partial reproductive isolation from each other due to different microhabitat and mating preferences (Rolán-Alvarez, 2007; Johannesson et al., 2010; Rolán-Alvarez et al., 2015a). This system is of particular interest in the Galician populations where these ecotypes live at different shore levels, while at the mid shore they meet and partially hybridize producing a hybrid zone that has been suggested as particularly suitable for ecological speciation in presence of gene flow (Rolán-Alvarez, 2007; Butlin et al., 2014; Boulding et al., 2017). Despite the number of empirical studies that have estimated biological and ecological parameters for this model system (reviewed in Pérez-Figueroa et al., 2005; Johannesson et al., 2010; Rolán-Alvarez et al., 2015a), there are several questions that remain unanswered (see below). Here, we perform spatially explicit computer simulations for studying the *L. saxatilis* Galician system incorporating the evolution of mate choice and reproductive isolation and using the parameter information available from empirical studies.

Noteworthy, there were previous efforts of modelling another *L. saxatilis* system, the Swedish one (Sadedin et al., 2009; Perini et al., 2020) which has also two ecotypes although under a different spatial and microhabitat distribution (see next section). Thus, although building on the Swedish 2009 modelling attempt, our model and simulations have different demographic and evolutionary settings, and utilize a different mating preference function for a better matching of the Galician demographic and evolutionary conditions (Rolán-Alvarez, 2007; Carvajal-Rodríguez and Rolán-Alvarez, 2014; Rolán-Alvarez et al., 2015a). The effect of introducing a cost of choosiness that was not considered in previous simulation studies was also included in our simulations. However, to validate our program we also replicate the 2009 Swedish model (see Program validation section).

Therefore, the main objective of this paper is try to answer the following questions which have been explicitly studied for the Galician system, although they can be extended to similar ecological speciation scenarios. Thus, given the conditions outlined in the Galician *L. saxatilis* model system (see details in Model and Methods but briefly, low dispersal ability, disruptive ecological adaptation, a magic trait, and an area of sympatry) our questions are:

1. Can we expect to find locally adapted specialists in the presence of gene flow?
2. Can mate choice (choosiness) and so full reproductive isolation, be evolved as a side effect of ecological adaptation?
3. Can the intensity of mate choice be increased at the area where the ecotypes meet because of reinforcement or other related mechanisms?
4. What is the impact of considering a cost in mate choice in relation to patterns of natural and sexual selection observed in nature?

Following, we briefly review the *L. saxatilis* species as a model organisms, then we explain the simulation model adapted to the Galician *L. saxatilis* scenario to answer the questions mentioned above.

## Material and Methods

### Model Organism

*L. saxatilis* is a marine snail gastropod living on North Atlantic rocky shores. The species crawls directly on rocks feeding on microalgae, diatoms and detritus. The species is gonochoric, shows internal fertilization and its ovoviviparous, showing direct development, i.e. with the female having a brood pouch with dozens to hundreds of embryos that are born as miniature snails. Due to these characteristics *L. saxatilis* shows low dispersal ability which also facilitates its capability to adapt to local conditions, and so local ecotypes have been frequently described on different localities and regions (Reid, 1996; Rolán-Alvarez, 2007; Johannesson et al., 2010; Rolán-Alvarez et al., 2015a). In particular, there is a striking sympatric polymorphism on Galician rocky shores (NorthWest Spain), where two ecotypes adapted to two different shore levels and microhabitats, coexist partially in sympatry at the mid shore (i.e. can frequently meet and mate partially assortatively, Rolán-Alvarez, 2007). A polymorphism determined by similar ecological forces (wave exposition and crab predation) is known in Britain and Sweden although the ecotypes are rarely (Britain) or never (Sweden) found in sympatry there (Rolán-Alvarez et al., 2015a). In Galicia, an upper shore ecotype (named “Crab” in (Butlin et al., 2014)) lives on the barnacle belt, showing a larger and more robust shell, coloured with alternate ribs and black lines, that protects against crab predation (Rolán-Alvarez, 2007; Butlin et al., 2014; Boulding et al., 2017). This ecotype also shows a good resistance to desiccation, osmotic and sun stress. In addition, the lower shore ecotype (named “Wave” in (Butlin et al., 2014)) appears associated to the mussel belt, and has a smaller, softer and smoother shell, with bigger aperture to accommodate a massive muscular foot. This strong foot is necessary to maintain attached the snail to the substratum due to the strong waves that commonly impact at first instance on the lower shore. There are no crabs that can predate on this ecotype on the lower shore, and so, corresponding upper and lower shore microhabitats, represent a differential selective regime for the ecotype subpopulations. At the mid shore, where the above microhabitats overlap forming a patched distribution, both ecotypes meet and hybridize producing a number of intermediate morphological forms, although they show partial reproductive isolation with each ecotype mating preferentially with specimens of the same ecotype. These intermediate forms may usually resemble genetically to one or another ecotype though they occasionally can be truly intermediate genotypes (Galindo et al., 2013; Kess et al., 2018). The partial premating reproductive isolation seems to be linked to the size differences existing between these two ecotypes and could be a side effect of the size assortative mating typically observed in the species, being considered as a typical magic trait (Boulding et al., 2017; Fernández-Meirama et al., 2017a).

### Simulation Model

There are few simulation studies on the interaction of natural selection with gene flow and the evolution of local adaptation in *Littorina* (Boulding, 1990; Pérez-Figueroa et al., 2005; Sadedin et al., 2009; Westram et al., 2018; Perini et al., 2020) and only the one from Sadedin and co-workers incorporates the evolution of mate choice and reproductive isolation.

The following model is a modification of the Sadedin et al. (2009) one, attending to the specific parameter values and spatial structure of the Galician *L. saxatilis* compared to the Swedish case. In the Swedish case the local morphological and behavioural adaptation occurs within islands in the Swedish archipelago in a horizontal wave exposure gradient through cliff and boulder habitats interspersed along the shore (Sadedin et al., 2009). On the contrary, the Galician ecotypes vary along a vertical, within locality micro geographic wave exposure gradient with different spatially varying selection favouring large sizes in upper-shore (wave-sheltered) and small sizes in lower-shore (wave-exposed). Unlike the Swedish case, the Galician ecotypes jointly with the intermediate forms, can be observed at approximately similar frequencies at the mid shore (Rolán-Alvarez, 2007; Galindo et al., 2013; Kess and Boulding, 2019). Migration rates of these snails are relatively low (less than 2 meters from the released point after one month), and the migration vectors of transplanted snails show a significant direction towards their native tidal height (Erlandsson et al., 1998).

The Galician simulation model begun with a pregnant female well adapted to the sheltered habitat arriving at the most upper deme in the shore, which is a realistic scenario of how *L*.*saxatilis* colonize new habitats (Janson, 1987). This founder female produced offspring having the optimum phenotype for the wave-sheltered shore and a random mating value for the mating trait (see below). The model was spatially explicit in one dimension with three shore levels (upper, intermediate and lower). Individuals were diploids with separate sexes, each individual was constituted by two different additive quantitative traits and a set of 8 microsatellite loci. The first quantitative trait is an ecological magic trait *x* i.e., it defines the individual fitness in a specific shore level and also is the target trait for the mate choice when mating is not random. An example of this kind of trait in *L. saxatilis* could be size (Boulding et al., 2017). The second trait *c* is a mate choice trait and is related with the sign of the preference for mate choice (positive or negative assortative) and also with the male choosiness. The choosiness is defined as the absolute value of the linear function *C* = 2*c* -1 while the sign of the preference depends on the sign of *C*. If *C* = 0 (*c* = 0.5) there is random mating; negative values of *C* (*c* <0.5) imply negative assortative mating, while positive values of *C* (*c* > 0.5) generate positive assortative mating (the absolute value, choosiness, varies between 0 and 1).

Each quantitative trait (*x* and *c*) was constituted by *L*∈{4,8} additive unlinked bi-allelic loci (see Table 1). Alleles may have value of 0 or 1. The trait value is scaled between 0 and 1 by summing over the alleles and dividing by *L*. Because we did not consider environmental variation the genotypic and phenotypic values were the same.

**Table 1.** Model parameters.

|  | Parameter | Symbol | Value |
| --- | --- | --- | --- |
| Demography | Generation number | $T$ | 20,000 |
| | Carrying capacity exposed deme | $K$ | 15,000 |
|  | Intermediate and sheltered demes |  | 3750 |
| | Mean offspring number | $b$ | 50 |
| | Between deme migration probability | $m$ | |
|  |  | 0 demes | 0.75 |
|  |  | 1 demes | 0.15 |
|  |  | 2 demes | 0.1 |
| Genome Structure | Number of microsatellites | | $L = 8$ |
| | Ecological magic trait ($L$ loci) | $x$ | $L = \{4, 8\}$ |
| | Male mating trait ($L$ loci) | $c$ | $L = \{4, 8\}$ |
| Selection | Selection strength | $\sigma_s$ | $\{0.15, 0.45, 1\}$ |
| | Selection in middle habitat | $\theta$ | $\{\text{no}, \text{yes}\}$ |
| Mating | Evaluated females per male | $N_f$ | 10 |
| | Mating preference tolerance | $\sigma_a$ | $\{0.05, 0.1\}$ |
| | Mate choice cost | | $\{\text{no}, \text{yes}\}$ |
| Mutation | Neutral mutation rate per locus | $\mu_0$ | $10^{-3}$ |
| | Trait mutation rate per locus | $\mu$ | $10^{-5}$ |

The generations were discrete non-overlapping and the simulation events occurred in the following order: birth, viability, mating, migration, reproduction and mutation. The parameter values can be consulted in Table 1.

### Spatial distribution

The spatial distribution of individuals in the simulation tried to resemble the Galician rocky shore where *L. saxatilis* inhabits (Rolán-Alvarez, 2007). The population was simulated as a one-dimensional array with a total of 26 demes. The first 20 demes corresponded to the upper shore which is the sheltered habitat in the Galician coast, the next two demes were the intermediate habitat and the last four corresponded to the lower shore which is the exposed habitat. Each deme had a maximum capacity depending on the carrying capacity of its habitat. A deme in the lower-shore exposed habitat had a carrying capacity of *K* = 15,000 while the intermediate and sheltered had *K* = 3750 each. This distribution of demes per habitat tries to match the true habitat distribution and densities observed in Galicia (Rolán-Alvarez, 2007). However, symmetric models (with equal number of demes and population density) were also run in order to interpret the consequence of this asymmetry (see below). Only pregnant females were considered as migrants. A migrant female can move through one or two demes, the offspring will be born at the arrival deme. Each individual develops its whole life within a deme.

### Viability

The survival condition of each individual depended on the distance between its ecological trait *x* and the optimum in the habitat where it was born. The optimum was 0, 0.5 and 1 for the exposed, intermediate, and the sheltered shore respectively. We also considered the case of no selection in the intermediate habitat so that under this scenario a fraction of individuals may reproduce in a spatial region (two demes) that is neutral with respect to ecological speciation (Cotto and Servedio, 2017). The fitness *w*_i_ of an individual *i* with phenotype *x*_i_ was given by (Sadedin et al., 2009)

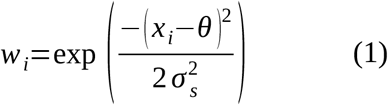

where *x*_i_ is the value of the ecological trait, θ is the optimum in the habitat where the individual was born and σ_s_ is the inverse of selection strength.

The offspring survival was density-dependent following a Beverton-Holt model (Beverton and Holt, 1957). Thus, the individual viability depended on the total number of local juveniles *N*, the mean number (*b* = 50) of offspring of each individual and the carrying capacity (Sadedin et al., 2009)

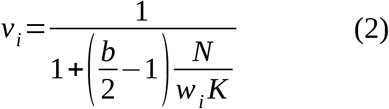

where *w*_i_ is the individual fitness (equation 1) and *K* is the carrying capacity of the deme.

### Mating

In *L. saxatilis* mate choice is performed by males (Rolán-Alvarez et al., 2015a). In our simplest model (without cost), each male always mate with one of 10 females randomly chosen from its own deme. The probability of mating between the male and any of the 10 females is proportional to the FND ψ function (*Ψ*_*FND*_, see equation 3).

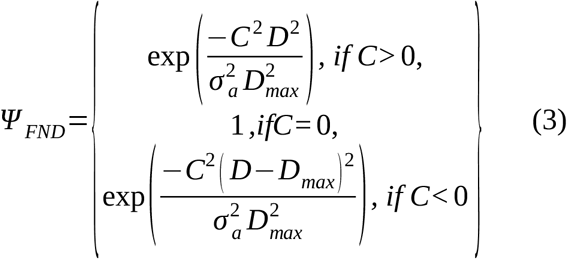

*Ψ*_*FND*_ depends on the value of the male mating trait *c* and the distance *D* = |*x*_m_ - *x*_f_| between the ecological traits of the male and the female candidate. The parameter *C* is a linear function of *c, C=* 2*c* - 1, and *D*_*max*_ is the maximum *D* (1 in this case) and *σ*_*a*_ is the tolerance or inverse strength of the male mating preference (Carvajal-Rodríguez and Rolán-Alvarez, 2014).

From (3), when *c* > 0.5 (*C* > 0) the mating is positive assortative, when *c* < 0.5 (*C* < 0) the mating is negative assortative, and it is random when the phenotype *c* = 0.5 (*C* = 0). The formulae in (3) is a modification of the original model of (Sadedin et al., 2009) to correct for the known problem that, when phenotypic values are equal to 0.5, positive assortative mating occurs regardless of the choice value (Carvajal-Rodríguez and Rolán-Alvarez, 2014).

As already explained, the choosiness is the absolute value of *C* while the sign of the preference depends on the sign of *C*. Thus, provided that some choosiness value evolved (|*C*| > 0), the resulting mate choice mechanism corresponds to a matching rule model combined with a magic trait (Kopp et al., 2018).

Under the mating without cost, each male always mates and produces offspring, independently of how choosy he is, so there will be mating even if all *Ψ*_*FND*_ values are very low. This is achieved by defining the mating probability as the FND function for a given male and female pair relative to the total FND function for all females that the given male encounters.

The following is an example of how mating is implemented without cost. Consider a situation where a male *i* encounters female 1 and they have *Ψ*_*FNDi1*_ = 10^−3^, and for any of the remaining females in the deme the pair of male *i* with any female *j* ≠ 1 has *Ψ*_*FNDij*_ = 10^−4^. The sum of *Ψ*_*FND*_ values for this male with the ten females he encounters is *S* = 10^−3^ + 9 × 10^−4^ = 0.0019. In the model without cost, the probability of mating for male *i* is Pr(*i*×1) + 9Pr(*i*×*j*) = *Ψ*_*FNDi1*_/*S* + 9*Ψ*_*FNDij*_/*S* = *S*/*S* = 1 where we divided by *S* due to the condition that the male must mate. Thus, we see that the probability of male *i* mating with female 1 is approximately 0.53 (i.e. 10^−3^/*S*) which is one order of magnitude greater than with any other female, so it is proportional to the *Ψ*_*FND*_ values.

### Mate choice cost

Given the value *Ψ*_*FND*_ of the preference function we considered also the case with cost to choosiness (Bolnick, 2004). In the model with cost, the mating probability is equal to the FND function for a given male and female pair (the coefficient of proportionality is set to 1) and therefore an overly choosy male might not find a mate. Following our previous example, male *i* randomly encounter female 1 plus 9 females from its deme and will try to mate with all of them but now the mating probability coincides with the *Ψ*_*FND*_ value and so, the probability of mating for male *i* is Pr(*i*×1) + 9Pr(*i*×*j*_≠1_) = *Ψ*_*FNDi1*_ + 9*Ψ*_*FNDij*_ = *S* = 0.0019. Therefore, there is more than a 99.8% chance that this male will not mate. If no mating occurs the male is discarded and next male is considered. If no mating takes place at all, the deme will be empty.

### Migration

The life cycle occurs within the deme and ends after the reproduction. The pregnant females can disperse 0 (no migration), 1 or 2 demes in either direction with probability 75%, 15% and 10% respectively. After migration, the female produce the offspring within the new deme and dies. These numbers were chosen in order to simulate a scenario of high gene flow under sympatry, as this was probably the starting point of the system before any local adaptation was evolved (Rolán-Alvarez, 2007).

### Reproduction

After mating and migration, the number of offspring was obtained from a Poisson with mean *b* (*b* = 50 see Sadedin et al., 2009). Recall that males are the choosy sex so each male randomly encounter some females in the deme and successful mating will occur depending on the FND function i.e. on the phenotypes of male and female and on the choosiness of the male. If a specific mating is successful then the offspring number per mated male-female pair is sampled from Poisson distribution with mean 50. The genetic composition of the offspring depended on the independent segregation of the trait loci and microsatellites from the parental gametes.

### Mutation

The new born underwent mutation after reproduction. The per individual haploid locus mutation rate was 10^−3^ for the stepwise mutation model of microsatellites (Slatkin, 1995) and 10^−5^ for the additive locus (Sadedin et al., 2009).

The life cycle was allowed to run during 20,000 generations. We performed 20 replicates for each parameter combination and 48 different cases of parameter combinations for the Galician model. The number of runs for some cases was extended to 80,000 generations to evaluate long-term results, as this number of generations corresponds to an upper threshold for Galician *L*.*saxatilis* based on a generation time of 6 months and estimates of divergence time between ecotypes (Quesada et al., 2007).

### Initial conditions

As already indicated in the simulation settings, the experiment begun with a founder event in which the most upper deme of the sheltered habitat was colonized by the offspring of a single female adapted to this habitat (*x* = 1). The offspring number was 50 (initial population size). All the individuals were homozygous for the quantitative traits and had the optimal ecological phenotype for this habitat (*x* = 1). The male mating trait was initially fixed at *c* = 0.5 (random mating). The microsatellite genotype was heterozygote for every locus and individual.

### Program validation

To validate our implementation we replicated two other ecological speciation simulation models, namely, the model in (Sadedin et al., 2009) concerning the Swedish *L. saxatilis* ecotypes and the model in (Gavrilets et al., 2007) concerning the speciation process of cichlids in a crater lake (Barluenga et al., 2006). This replication effort comes in the spirit of exploring the robustness and reproducibility of individual-based models in order to augment credibility and consistency of computational modelling and simulation and thereby improving the evolutionary ecology theoretical field in general and speciation in particular (Massol and Debarre, 2015; Thiele and Grimm, 2015; David et al., 2017).

Regarding the replicated models, we have implemented only the mating system by similarity (matching rules sensu Kopp et al., 2018) with asymmetric resource availability. Also, regarding the cichlids model we have implemented only the two patches space structure. The replication of such models allowed us to validate our implementation while at the same time it permitted to interpret the sensitivity of the evolutionary outcome to some of the assumptions in the model, as the type of preference function and the presence or absence of mate choice cost.

### Analysis

We analyse the data obtained in the simulations using different variables. For example, the occurrence of ecological adaptation to the exposed habitat (lower shore) was recorded when the ecological trait changed its mean value by more than 75% (*x* < 0.25) as it is an arbitrary but reasonable indicator that most individuals have adapted to the new habitat and similar thresholds have been used in the past (Sadedin et al., 2009).Similarly, reproductive isolation evolution was recorded when the mean choosiness value was higher than 0.1 or, that is the same, the mean value of *c* was appreciably different from 0.5. (*c* < 0.45 or *c* > 0.55). In addition, we measured the genetic differentiation between habitats by means of the *F*_ST_ (Wright, 1951) and *Q*_ST_ indices (Leinonen et al., 2013). The fixation index *F*_ST_ was computed for the microsatellites as *F*_ST_= 1- (((*H*_low_ + *H*_mid_ + *H*_up_)/3) / H_total_) where *H*_shore_ was the heterozygosity at the lower, middle or upper shore levels, and *H*_total_ was the total heterozygosity without considering the shore levels. Similarly, the quantitative genetic analogue of *F*_ST_ was computed for the ecological trait as *Q*_ST_= *V*_between_ / (*V*_between_ + 2\**V*_within_) where *V* is the variance between or within habitats. Hence, if high mating discrimination happened in the middle habitat, *c* ≥ 0.9, jointly with the occurrence of ecological adaptation, *Q*_ST_ ≥ 0.9, then this was considered as reproductive isolation between the two ecotypes and can be interpreted as a positive case of speciation caused by ecologically-based divergent natural selection i.e. ecological speciation.

Simulation studies should be statistically analysed in order to be properly understood and gain scientific credibility (Lotterhos et al., 2018). To evaluate the different simulated scenarios in the Galician model, we investigated various simulation outcomes (dependent variables) that could be influenced by *a priori* different combination of simulation parameters (independent variables) under a classical multifactorial ANOVA (see studied variables in Table 2). First, our independent variables were combined using an orthogonal design (24 parameter combinations x 20 replicates per combination).

**Table 2.** Independent (factors) and dependent (response) variables for the ANOVA.

|  |  |
| --- | --- |
| **Independent variables** | Selection strength ($\sigma_s$) |
| | Mating strength ($\sigma_a$) |
| | Number of loci ($L$) |
| | Selection in the mid-shore ($\theta$) |
| **Dependent variables** | Ecological trait ($x$) |
| | mating trait ($c$) |
| | Mating correlation for $x$ ($r$) |
| | Population size ($N$) |
| | Differentiation measures ($F_{ST}$ and $Q_{ST}$) |

Notice that we do not use ANOVA as classical way for hypothesis testing, but rather as an exploratory tool to compare the relative contribution of different factors (and interactions) into the dependent ones. The relative contribution of each factor to the overall variation was measured by the eta-squared coefficient (Sokal and Rohlf, 1981). The eta-squared is finally expressed as % with respect to the sum of all etas in the ANOVA (in order to allow comparison among different ANOVAs). In this way we can identify the most interesting relationships (those with higher eta-squared i.e. higher proportion of variance of the dependent variable explained) which can be detailed *a posteriori* in the following sections. The per habitat average of the quantitative trait phenotypic values, *x* and *c*, were used as dependent variables under this ANOVA approach, and also the within habitat population size (*N*, considering all demes per habitat), as well as the *F*_ST_ and *Q*_ST_ estimates described above. Additionally, we used the Pearson correlation (*r*) of *x* values among partners in mates within habitat, as a way of estimate the assortative mating for the ecological (magic) trait.

## Results

### Program validation

When developing our Galician model we adapted it to replicate the 2 × 1 model (two habitats, 1 deme per habitat) in (Gavrilets et al., 2007) concerning the speciation process of cichlids in a crater lake (Barluenga et al., 2006). We obtained the same results regarding speciation events, choice trait and *F*_ST_ values, as reported from the original article. We also reproduced the similarity scenario for the Swedish *L. saxatilis* model *(Sadedin et al*., *2009)* using their same mating preference function and obtained qualitatively the same results regarding adaptation, neutral genetic differentiation, ecotype formation and non-random mating evolution.

Finally, we repeated the simulations with the Swedish model but using the preference function defined in (3) instead of the original one in (Sadedin et al., 2009). We obtained some new cases of evolution of negative assortative mating and fewer invasion and adaptation cases. These results are expected since it is known that the original function in (Sadedin et al., 2009) has an anomalous negative assortative mating behaviour (Debarre, 2012) implying less negative assortative mating cases than expected (Carvajal-Rodríguez and Rolán-Alvarez, 2014). However, the main conclusions in Sadedin remained, namely, ecotypes may persist indefinitely with moderate genetic differentiation without speciation. As Sadedin *et al* highlights, both ecological divergence and the evolution of reproductive isolation, are crucial to ecological speciation, but since they are driven by different forces that can even conflict, the prediction about ecological speciation requires highly detailed data jointly with a quantitative simulation model specially designed for the problem at hand as we did with what we have called the Galician model.

### Galician model

The results are first explored by factorial ANOVA, in order to emphasize the main factors (and interactions) contributing the most (highest variance explained) to the dependent variables, and secondly the details of these effects are fully described in next sections.

### Parameter effects: Orthogonal analysis of variance

The relative importance of the different simulated parameters, as depicted by the selection and mating strength, number of genes, etc, was summarized by the ANOVA from which the factors and interactions that explained the larger percentage of variance in the evolutionary outcome can be outlined (Table 3, also see supplementary material for the full table of the factorial ANOVA).

**Table 3.**
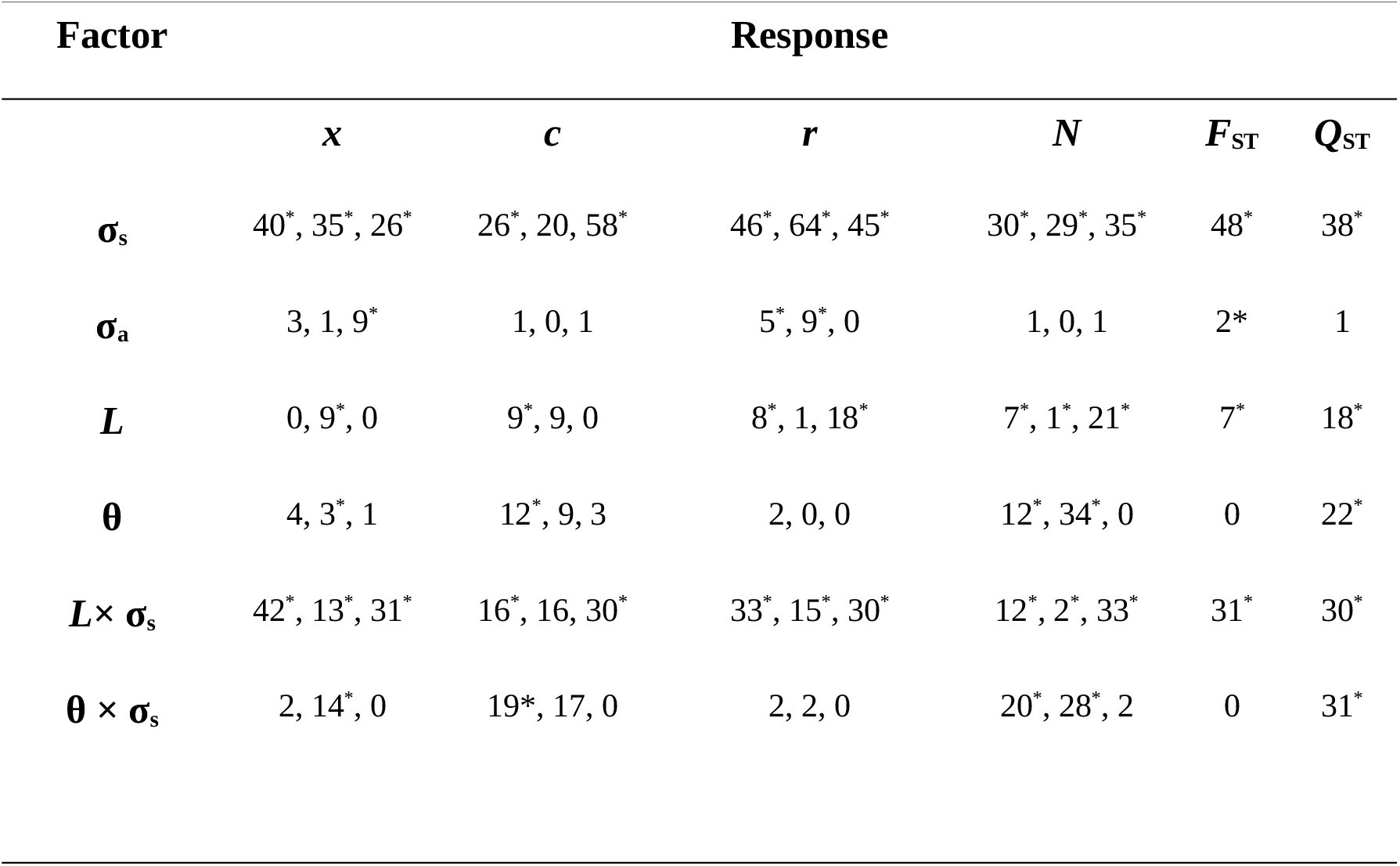
ANOVA results showing % of variance (% partial eta squared; see text) explained by each factor for the ecological (*x*) and mating (*c*) traits, mating correlation (*r*), population size (*N*) and differentiation measures (*F*_ST_ and *Q*_ST_). Only the two most important factor interactions are presented. The values within cells correspond to the % at lower, middle, upper shore, by this order, except for differentiation (*F*_ST_ and *Q*_ST_) that are between shore levels. The asterisk indicates significance at the 0.001 level.

| Factor | Response |  |  |  |  |  |
| --- | --- | --- | --- | --- | --- | --- |
| | $x$ | $c$ | $r$ | $N$ | $F_{ST}$ | $Q_{ST}$ |
| $\sigma_s$ | 40*, 35*, 26* | 26*, 20, 58* | 46*, 64*, 45* | 30*, 29*, 35* | 48* | 38* |
| $\sigma_a$ | 3, 1, 9* | 1, 0, 1 | 5*, 9*, 0 | 1, 0, 1 | 2* | 1 |
| $L$ | 0, 9*, 0 | 9*, 9, 0 | 8*, 1, 18* | 7*, 1*, 21* | 7* | 18* |
| $\theta$ | 4, 3*, 1 | 12*, 9, 3 | 2, 0, 0 | 12*, 34*, 0 | 0 | 22* |
| $L \times \sigma_s$ | 42*, 13*, 31* | 16*, 16, 30* | 33*, 15*, 30* | 12*, 2*, 33* | 31* | 30* |
| $\theta \times \sigma_s$ | 2, 14*, 0 | 19*, 17, 0 | 2, 2, 0 | 20*, 28*, 2 | 0 | 31* |

The selection strength σ_s_ was clearly the key element with highest effect in adaptation, choice evolution, population size and differentiation. The number of genes had typically a significant although lower impact on most dependent variables. The mating strength (σ_a_) and selection on the mid shore (θ) showed in general low impact on most variables. On the other hand, the interaction of selection with the number of genes, and with less importance the θ × σ_s_ interaction, had also great importance for most variables. Some factors require special attention in certain variables. For example, the number of genes (*L*) shows a moderate impact in the mating correlation (*r*) and both *F*_ST_ and *Q*_ST_ variables. In the following we present these outcomes more in detail.

### Ecotype formation: Habitat colonization and adaptation

The founder population had the ecological trait perfectly adapted (*x* = 1) to the sheltered (upper shore) habitat. As time passed some individuals migrated to the intermediate and lower habitats where the ecological conditions were different so that the optimal value for the exposed habitat (lower shore) was *x* = 0. In the previous section, we have emphasized that σ_s_ and *L* × σ_s_ were the most relevant factors determining the evolution of the ecological trait *x* (Table 3). We will now describe these patterns in further detail. In Figure 1 we may appreciate the summary of the adaptation process after 20,000 generations, through the interplay between migration and selection within habitat, averaged over the different simulated scenarios and demes.

**Fig 1.**
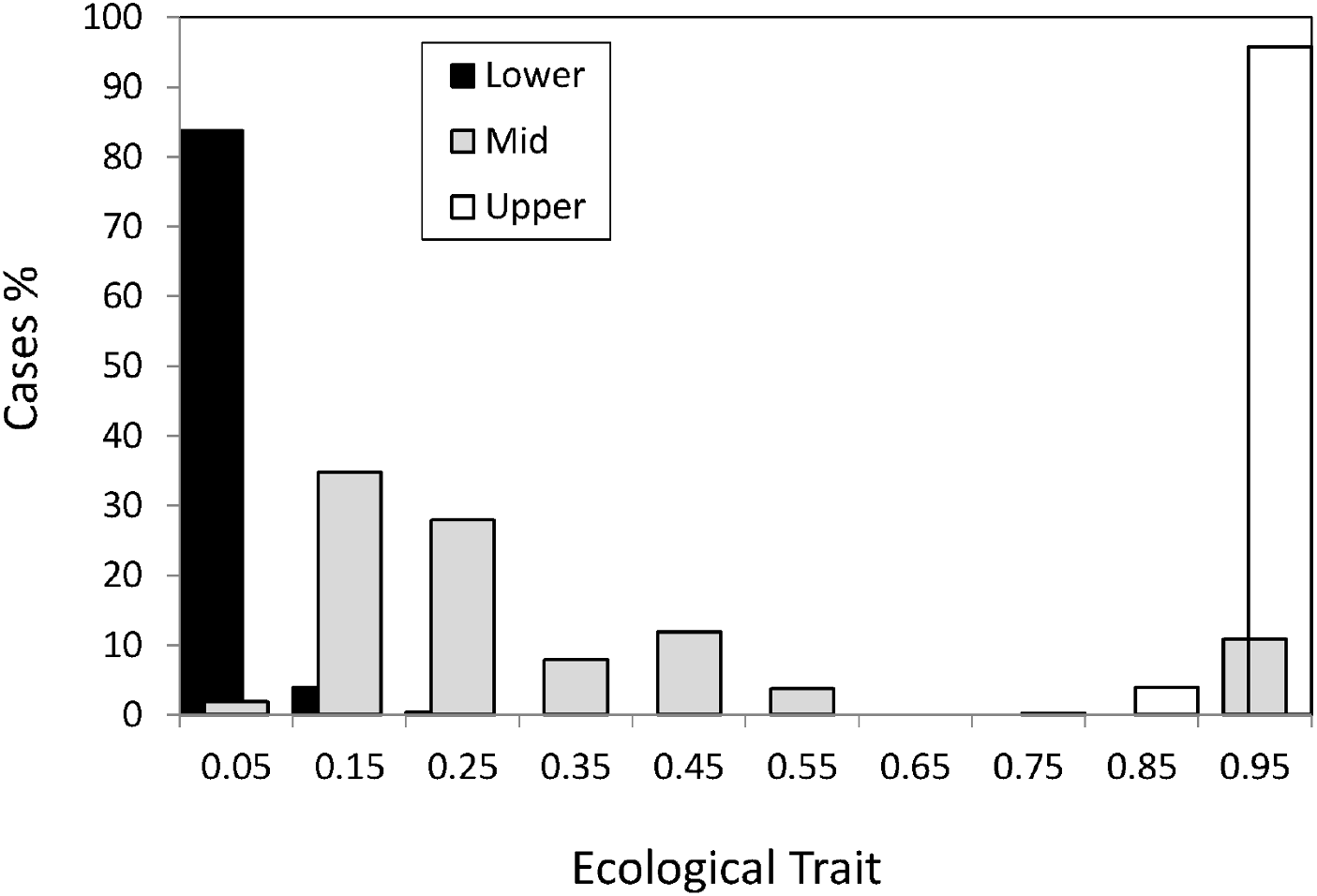
Mean value per habitat of the ecological trait for individuals living in the sheltered (upper), intermediate and exposed (lower) habitats.

First, adaptation to the upper shore (Figure 1, white bars, *x ≥* 0.75) happened in virtually all cases which is not surprising since the founder individual was perfectly adapted to this habitat.

Habitat colonization and adaptation to the lower shore happened under weak to intermediate selection (σ_s_ ∈ {1, 0.45}), irrespective of the value of the other parameters. Under strong selection (σ_s_ = 0.15) adaptation occurred in cases with few loci (*L*=4) and non-neutral hybrid zone (optimum 0.5) and also under neutral hybrid zone and mating strength σ_a_ =0.1. For cases corresponding to scenarios under strong selection and more number of loci (*L*=8) adaptation only occurred when the hybrid zone was non-neutral. Thus, we may appreciate that more than 80% of the simulated cases successfully colonized the new habitat and adapted to it having the optimal ecological phenotype (Figure 1, black bars, *x* ≤ 0.15, Fig. 1).

The middle shore (Figure 1, grey bar), had some cases where generalists (0.4 < *x* < 0.6) evolved. The evolution of generalists in the middle shore corresponded to scenarios with non-neutral hybrid zone and strong selection (σ_s_ = 0.15) with adaptation in the lower shore. Other cases under strong selection evolved ecological trait phenotypes close to the upper shore optimum (Figure 1, *x* > 0.75). In the latter cases, there was not colonization of the lower shore and they only differ from cases in which generalists evolved in that the ecological trait was neutral in the hybrid zone. Thus, under strong selection in the upper and lower shores, the environmental conditions of the hybrid zone as defined by the selection in the mid-shore (θ) are key to the evolution of generalists and the colonization of the lower shore.

However, more than 60% of the cases in the middle habitat had an ecological trait value closer to the optimum for the lower habitat (Figure 1, *x* < 0.25). This asymmetry, i.e. ecological trait values in the intermediate habitat being closer to the lower shore than to the upper shore optimum, can be explained by the higher carrying capacity of the lower habitat which implied a higher gene flow from lower to middle than from upper to middle through the vertical axis of the Galician coast.

To confirm the cause of the asymmetric effect for adaptation in the middle habitat, we studied a specific symmetric case corresponding to *L* = 4 loci under intermediate selection (σ_s_ = 0.45) and neutral hybrid zone. We simulate 100 replicates of the Galician model and also 100 replicates of a symmetric model with the same number of demes in the upper and lower shore and equal per deme carrying capacity in the three habitats (supplementary Table S1). We found no differences in adaptation to the lower shore and confirmed the effect of differential migration to adaptation to the middle shore where the mean ecological trait value was 0.15 ± 0.04 for the Galician model while it was 0.48 ± 0.07 for the Symmetric model (supplementary Table S2).

The asymmetric effect is a significant difference with the Swedish model where intermediate zones are non neutral (optimum 0.5) and their distribution is not localized just in the middle point of the vertical coast axis but in the circular horizontal borders between sheltered and exposed so before each sheltered and at the end of each exposed in a circular (as in an island coast) distribution.

In addition to the ecological trait values, we also studied the time to adaptation to the exposed habitat. Basically there were only two opposite outcomes, most scenarios produced fast adaptation in less than 1000 generations but some did not reach adaptation in 20,000 generations (they appear as the right-most bar in Fig.2). As before, the strength of selection was the fundamental factor affecting the speed of adaptation. Most fast adaptation cases involve an intermediate-low selection strength while most cases of non-adaptation corresponded to strong selection.

**Fig 2.**
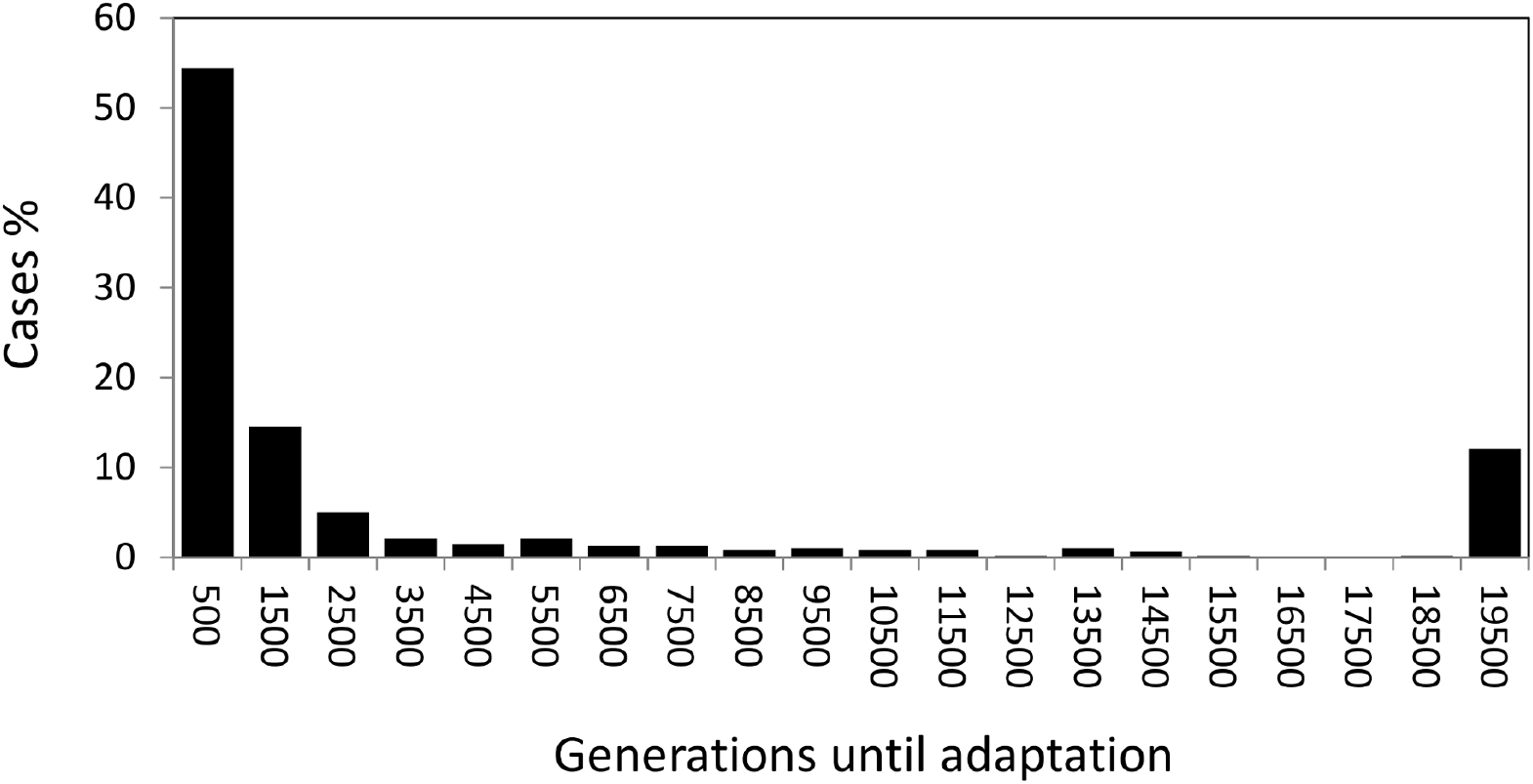
Generations until adaptation to the lower shore (*X* < 0.25) for the different scenarios assayed.

There was also an interaction effect between the strength of selection and the number of loci (*L* × σ_s_ in Table 3), implying that under strongest selection (σ_s_ = 0.15) there was twice the probability of adaptation under the fewer loci scenario (*L* = 4) than under the greater one (*L*=8).

### Mating trait

We have already emphasized that it was σ_s_ and σ_s_ × *L* the most consistent factors determining the evolution of *c* (Table 3). We will now describe how these effects occur in detail. The phenotypic value for the male mating trait, *c*, was initially 0.5 that implies random mating (i.e. no choosiness |*C*| = |2*c* -1| = 0). Values of *c* above 0.5 imply positive assortative while below 0.5 imply negative assortative mating. To discard noisy variation around 0.5 we considered that positive mate choice evolved when the mean *c* values were above 0.55 while negative choice evolved when mean values were below 0.45. Thus, from Fig. 3 we appreciate that positive assortative mate choice (*c* > 0.55) evolved in the three habitats in most cases although there were some cases (about 15% in the three habitats) with negative mate choice (*c* < 0.45, Fig. 3).

**Fig 3.**
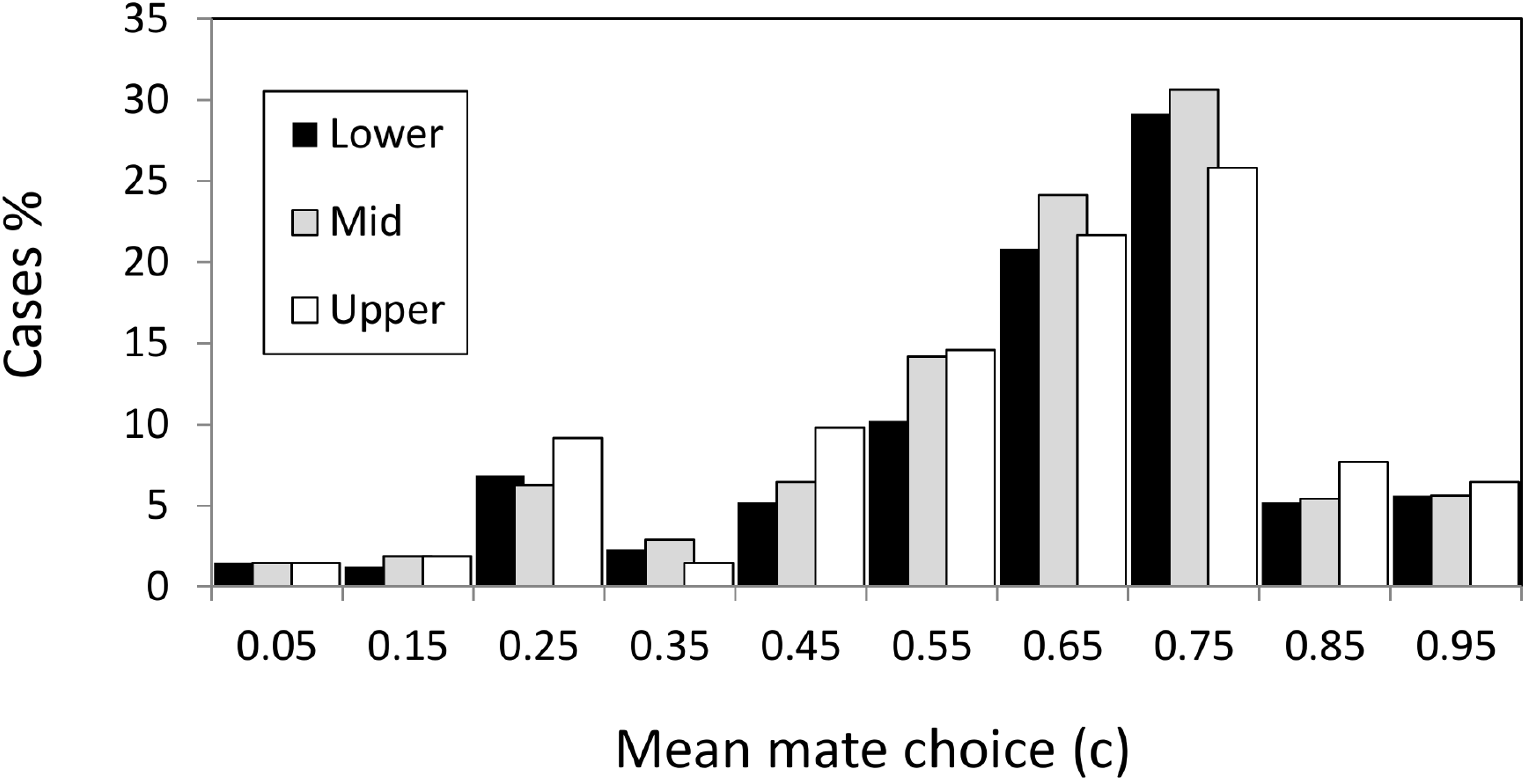
Mean value of the mate choice trait for individuals living in the upper (sheltered), middled and lower (exposed) habitats.

There were a clear pattern behind the negative assortative mating scenarios, the combination of few loci (*L* = 4) with strong selection (σ_s_ = 0.15) (Table S3). Negative assortative mating only evolved under strong selection scenarios and the most favourable scenarios include few loci, low tolerance (σ_a_ = 0.05) and the selective middle zone. It seems that under this scenario it was useful to evolve a preference for the different type in order to maintain the polymorphisms avoiding the fast fixation of sub-optimal genotype combinations. If fact, these negative assortative mating cases did not evolve fast, i.e. it took at least 1500-10,000 generations until the *x* trait was below 0.25 in the exposed shore and after 20,000 generations the mean fitness still was sub-optimal (*w* < 0.9) in the three habitats.

When selection was not so strong and/or the number of loci was high, the result was mostly positive assortative mating (70% of cases in the three habitats with average choosiness of 0.45, *c* = 0.725, see next section).

As with the ecological trait, we also studied the time required to evolve choice in the lower habitat. Interestingly, the time needed for this character to evolve is much longer than for the ecological trait.

### Mating trait *versus* mating correlation

The increase of reproductive isolation in speciation scenarios is typically estimated by means of size (or ecologically related traits) assortative mating (Jiang et al., 2013; Janicke et al., 2019). However, the evolution of assortative mating can be only produced by means of the mate choice (choosiness in our model). Therefore, it is of special interest to compare a genetic trait that has evolved, such as male mating trait (*c*) values, with the observed mating pattern, as measured by the correlation (*r*) between male and female phenotypes *x* (ecological trait) in mated pairs, since this is a common measure of assortative mating (Fig. 4). This could help to identify potential situations in which demographic effects may give the wrong impression of mate choice evolution.

**Fig 4.**
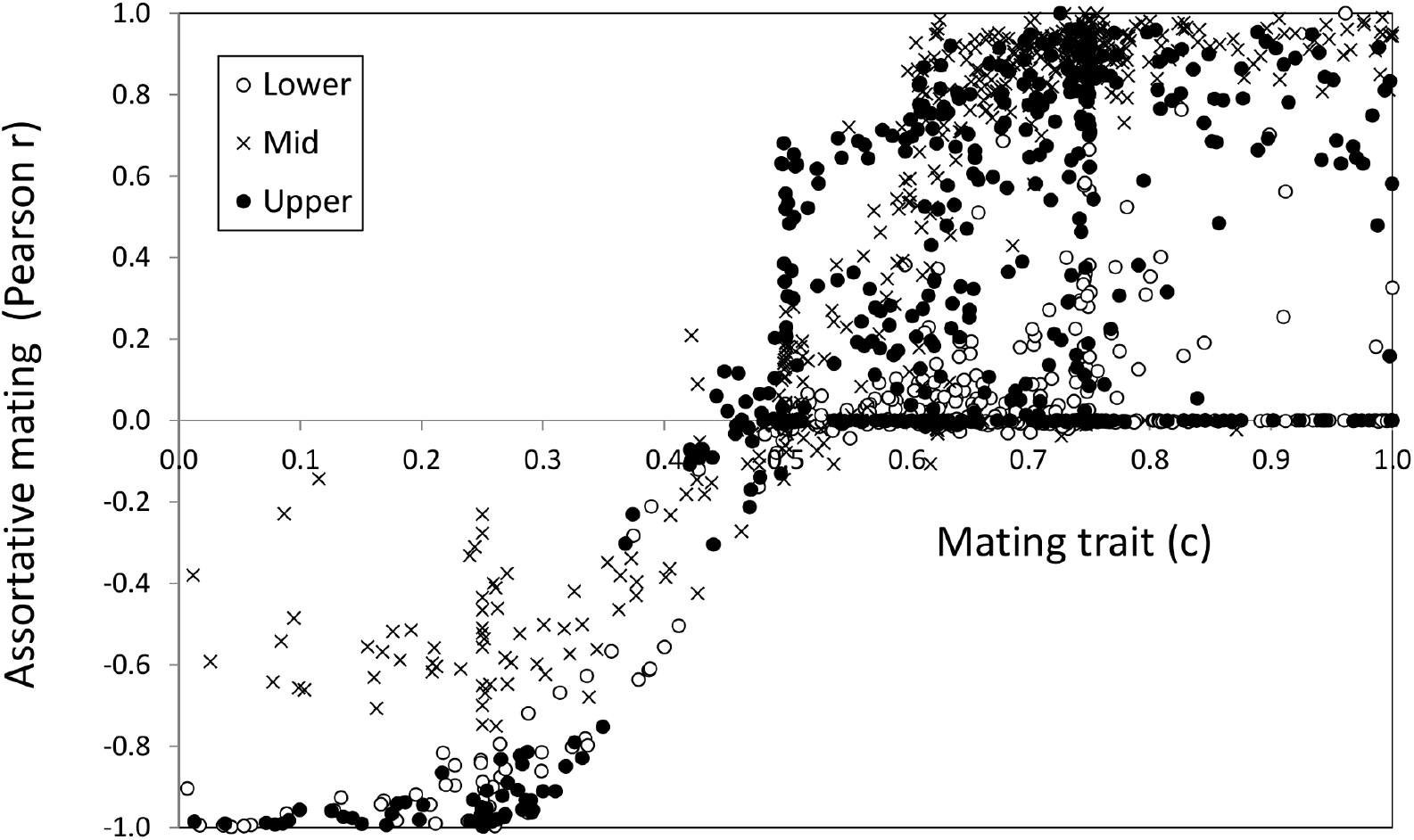
Mating trait value (*c*, abscissa axis) versus the mating pattern as measured by the correlation (*r*, ordinate axis) for the ecological trait in mated pairs.

There was good agreement between the sign of the choice and the mating pattern described by the correlation. Regarding choosiness, higher |*C*| values implied higher absolute correlation value and vice versa. The agreement was quite good in the range 0.2 < *c* < 0.8 i.e. a choosiness below 0.6 (|*C*|< 0.6). With choosiness values above 0.6 (*c* ≤ 0.2 and *c* ≥ 0.8 values) there was a saturation of the correlation value reaching its maximum in several cases. Also noteworthy is that there were some cases with high choosiness that presented very low correlation (points close to the abscissa at the right-hand end of Fig. 4). These latter cases corresponded to scenarios where the ecological trait was virtually fixed.

In general, about 1/3 of the cases, (33%, 34% and 29% for the lower, middle and upper shore respectively) reached intermediate (0.4-0.6) positive choosiness values as a result of the trade-off between divergence and sexual selection thus favouring local adaptation under migration (Rettelbach et al., 2013; Cotto and Servedio, 2017; Sachdeva and Barton, 2017).

As with the ecological trait, we studied the possible effect of the asymmetry of the Galician model in the evolution of the mating trait and observed that in this case the asymmetry in carrying capacity seems to have little or no effect on the evolution of the mating trait (supplementary Table S2). The values obtained for the mating trait matches well the empirical estimation of mate choice values for *L. saxatilis* and other littorinids (Fernández-Meirama et al., 2017a, 2017b).

### Neutral (*F*_ST_) and quantitative (*Q*_ST_) genetic differentiation

Regarding the relative contribution of the distinct factors to the population differentiation, the analysis of variance (Table 3) showed that the selection strength (σ_s_) and its interaction with number of loci (*L*×σ_s_) had the highest effect on both neutral and quantitative differentiation. In addition, all interactions in the table involving strength of selection (σ_s_) as well as the number of loci (*L*) showed a large impact on quantitative differentiation (*Q*_ST_). This suggest the great importance of the strength of selection (as a factor and in interactions) in determining the variance explained in Q_ST_. However, the ANOVA did not investigate the relationship between *F*_ST_ and *Q*_ST_.

The *Q*_ST_ *vs F*_ST_ comparison (Fig. 5) presented the typical pattern of local adaptation in the presence of gene flow, with high *Q*_ST_ versus low *F*_ST_ values (Leinonen et al., 2013). The only exception occurs when adaptation is not reached within the 20,000 generation interval (e.g. strong selection with *L*=8 loci and neutral hybrid zone, see the ecotype formation section), in that cases the *Q*_ST_ values remained low.

**Fig 5.**
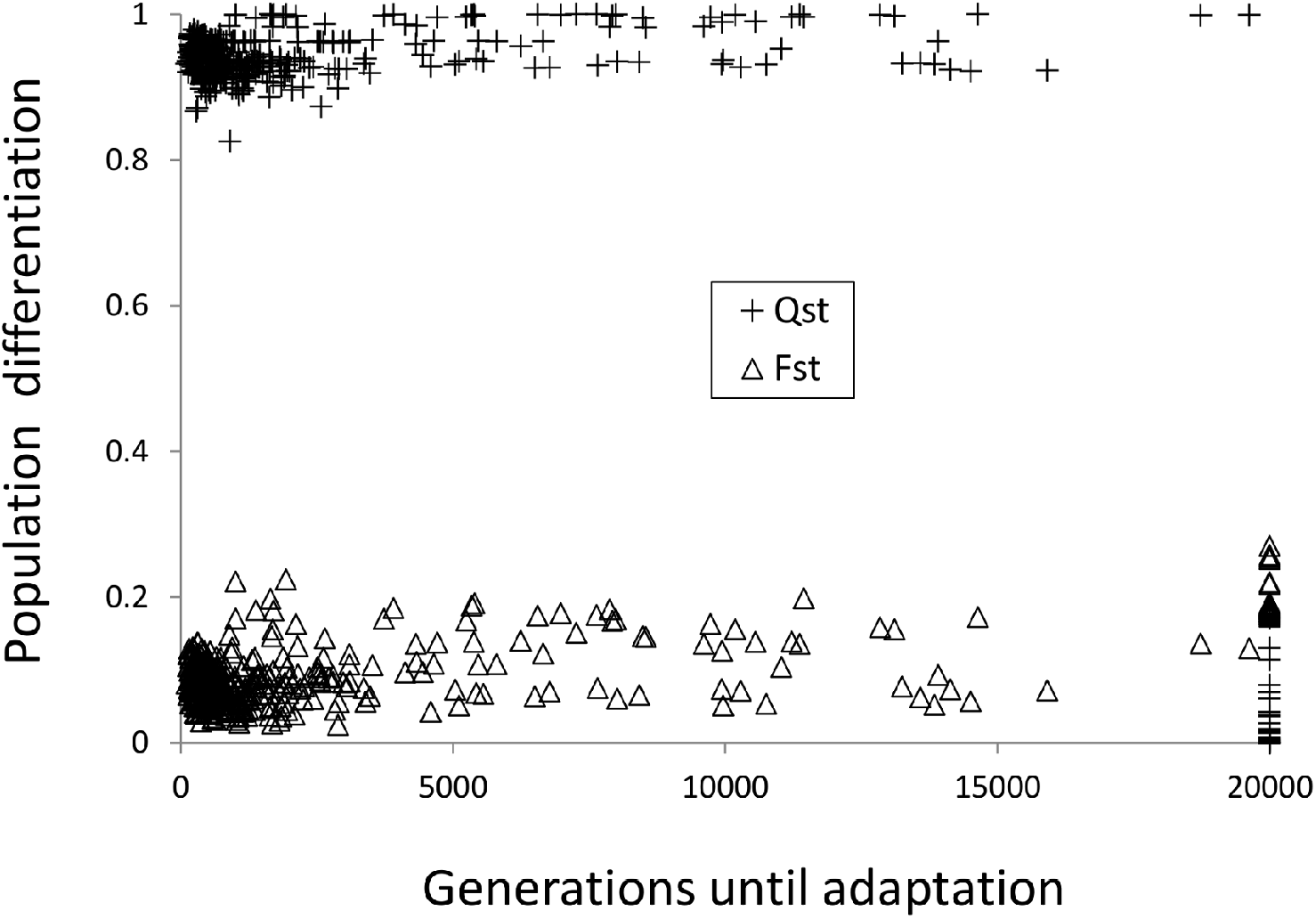
Neutral versus quantitative genetic differentiation

### Intermediate habitat effect

We have modelled two different ecological scenarios for the intermediate habitat (2 demes). In the selective-middle habitat scenario there is positive selection for the ecological trait with the optimum in 0.5. In the neutral-middle habitat scenario the ecological trait was neutral. The latter is a realistic scenario not usually considered in ecological speciation modelling studies (but see Cotto and Servedio, 2017). The kind of scenario (selective or not) had a minor impact (affecting mean trait values) regarding the ecological trait (*x*) in the middle shore but a moderate to high impact for *c* (at lower shore), *N* (from low to mid shore) and *Q*_*ST*_ (Table 3).

Both under the selective and neutral-middle scenarios, there were in the middle shore, a higher percentage of cases with mean ecological phenotype closer to the lower zone optimum (47 or 64% for selective or neutral-middle respectively) than to the upper zone optimum (0 or 11% for selective or neutral-middle respectively). This asymmetry was already evident in Fig. 1, and was related with the Galician microhabitat configuration jointly with the higher carrying capacity of the lower habitat that provoked a higher number of migrants arriving from this habitat (Table S2). This result matches the empirical observation regarding the size of hybrids in the mid-shore which even though they are genetically more heterogeneous than Crab (upper shore) or Wave (lower shore) ecotypes they tend to be closer to the lower shore phenotype (Galindo et al., 2013; Diz et al., 2021).

Regarding the impact of the middle scenario over the colonization of the lower shore, there were more colonization cases of the exposed habitat when there was selection in the middle shore (97% of those cases) than when the middle shore was neutral (79% of those cases). In fact the analysis of variance (Table 3) showed that the selection strength (σ_s_) and its interaction with the middle scenario (θ × σ_s_) had significant effect for the *x* trait in the middle zone.

Regarding the mating trait and mating correlation, the middle habitat scenario did not contribute to explain any significant variance for the mating trait or the correlation over the simulations (Table 3 and compare rows within Table S4 and Table S5). Also, the choosiness between the different habitats were quite similar (compare columns in Table S4). However there were important differences for the correlation between habitats. The mean mating correlation was 0.24 in the exposed area, 0.55 in the middle and 0.34 in the sheltered area. However, these general mean values are affected by cases with negative choice and also by cases with poor adaptation. Obviously, when excluding negative choice scenarios the correlation increases but the pattern with higher correlation in the middle habitat still holds. Specifically for the intermediate selection cases, the correlation values in the middle habitat can be above 0.85 (Table S5). The effect was even higher when the symmetric model is considered (Table S6) so that the higher correlation in the middle habitat was not an effect of the asymmetry in the Galician model.

Finally, regarding genetic differentiation, *F*_ST_ was not affected by the presence or absence of selection in the intermediate habitat, contrary to quantitative differentiation (*Q*_ST_) which was 20% higher in average when the middle habitat was selective (*Q*_ST_ = 0.93), compared to when the middle was neutral (*Q*_ST_ = 0.74). The type of middle habitat scenario explained 22% of the variance in *Q*_ST_ over the simulations (Table 3).

### Ecological speciation

We defined as a proxy for complete reproductive isolation the concurrence of high local adaptation (*Q*_ST_ ≥ 0.9, see Fig. 5) and high middle-shore choosiness (see Fig. 4, *c* ≥ 0.9 i.e. positive assortative mating with choosiness |*C*| ≥ 0.8). This reproductive isolation caused by ecologically-based divergent natural selection is called ecological speciation. Thus, when taking jointly the evolution of adaptation and assortative mating, we observed that there were ecological speciation in about 5% of the simulations. All the ecological speciation cases happened under the few selective loci (*L*=4) scenario. However, they were not uniformly distributed for the different selection strengths with all speciation cases linked to intermediate or weak selection strength. We increased the simulation runs to 100 to check this result and observed an 11-15% percentage of speciation for intermediate and weak selection while there were no speciation for the strong selection cases (σ_s_ =0.15, Table S7). These same results were also obtained in the long-term (80,000 generations).

### Mate choice cost

As before, the relative importance of the different evolutionary scenarios was summarized by the ANOVA (supplementary Table S8). The strength of selection was again the most important factor influencing the dependent variables. The factor *L*× σ_s_, i.e. the interaction between number of loci and strength of selection, had decreased importance. In general many more factors were now non significant in several variables, which suggest that with cost less adaptation and evolution of variables is obtained.

In the previous scenarios there were no cost for being choosy. Adding a cost to the mate choice implies that the overly choosy males may remain unmated. As a result there were no evolutionary outcome in the form of negative choice neither negative assortative mating (negative mating correlation). Also, less positive choice was evolved in the three habitats (see Table 4).

**Table 4.**
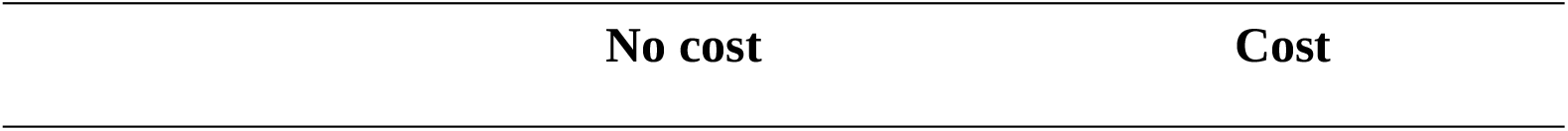
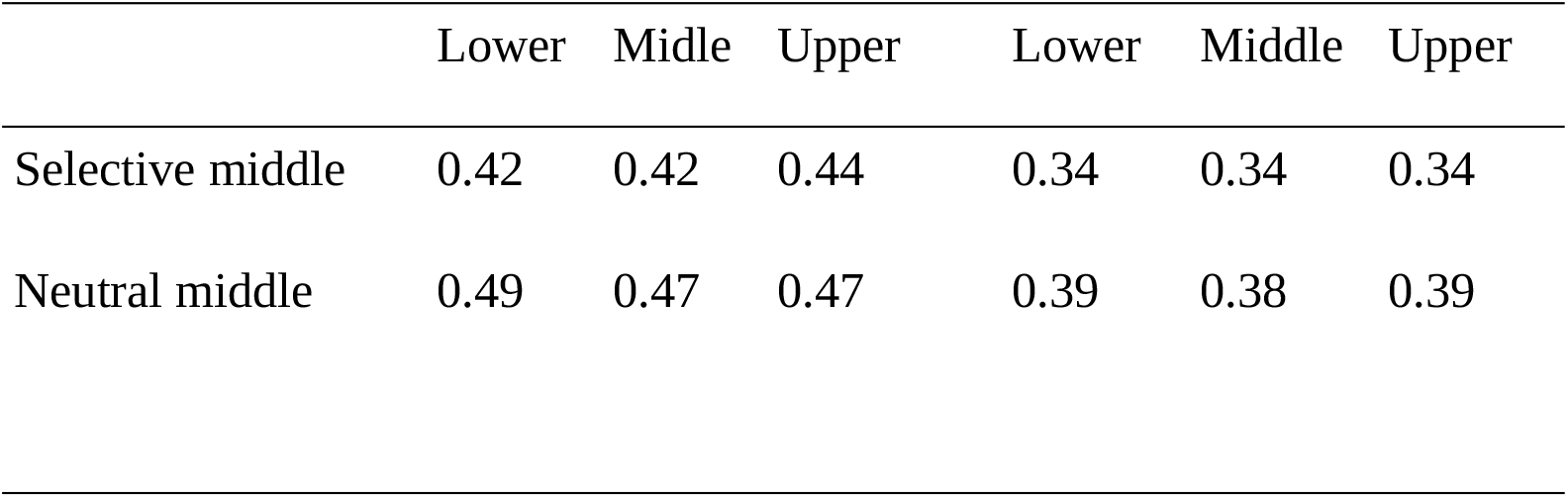
Mean choosiness |*C*| linked to positive assortative mating (negative values excluded) for the different habitats with selective or neutral middle scenario and with or without mate choice cost. The values are averages for the different *L* and σ_a_.

Finally, there were fewer colonization events in the presence of cost. Under the selective-middle habitat scenario 83% of the simulations with cost underwent colonization, compared to 97% without cost. The same pattern occurred under the neutral-middle habitat scenario, 66% of colonization with cost versus 79% without cost. However, when colonization occurred the fitness in the exposed habitat was higher under cost scenarios (mean fitness 99.7%) indicating that adaptation was better in these cases. Finally, we do not find ecological speciation when mate choice cost was included but in just one run of the case with weak selection (σ_s_ = 1), low number of loci (*L*=4), higher tolerance (σ_a_ = 0.1) and selective-middle scenario. This case implies a 0.2% compared to 5% speciation cases when cost was absent.

## Discussion

We have analysed the evolution of local adaptation and mate choice in a system displaying an extreme microhabitat environmental heterogeneity. This system resembles the *L. saxatilis* microhabitat-associated dimorphism along the wave-exposed rocky shores of Galicia. *L. saxatilis* dimorphism is a good example of microgeographic adaptation and incipient-speciation which occurs despite high gene flow based on the species expected levels of dispersal (Rolán-Alvarez 2007; Richardson et al., 2014). In our individual-based simulations we investigated the dynamics of the ecotype formation through the inter-deme and habitat spatial and temporal scales. This deme-based fine spatial scale model showed how the strength of selection and the evolution of choosiness may interact in order to produce local adaptation and reproductive isolation (Cotto and Servedio, 2017).

Our results are consistent with the key role of natural selection in the ecological speciation processes (Barton, 2010). The strength of selection on its own or through interaction with other factors explains most of the variance in the dependent variables over the scenarios simulated for both with and without mate choice cost. That is, most of the differences in ecological and mating trait values are explained by the strength of selection.

We have seen how the population, as the individuals disperse through inter-habitat demes, evolve different phenotypic values for habitat specialization. Given the high environmental heterogeneity between the sheltered and exposed habitats we obtained specialists for these habitats. The outcome was slightly different in the middle habitat, where generalist or specialist evolved depending on how the environmental heterogeneity was defined. When the middle habitat was neutral this facilitated the co-existence of specialists from both shore levels although with a higher % from the lower shore due to the higher number of migrants coming from this shore. On the contrary, when the middle habitat was non-neutral with an optimum in-between both extreme shore levels, which implies a less-heterogeneous environment, more intermediate phenotypes evolved. These results are consistent with previous studies showing the positive relationship between local adaptation and environmental heterogeneity (Berdahl et al., 2015; Svardal et al., 2015).

Concerning our question number 1 about the pattern of local adaptation under gene flow, which expects high quantitative genetic divergence for certain traits, without a general divergence at the genome level, we confirmed the expected pattern and observed high quantitative trait genetic divergence (*Q*_ST)_ jointly with low neutral genetic differentiation (*F*_ST_).

We found that the time-scale for ecological adaptation and the occurrence of ecological speciation were influenced by the strength of selection. If selection was strong the time for adaptation was longer. Moreover, weak or intermediate selection and few selective loci were favoured under ecological speciation. However it should be noted that we modelled independent loci, i.e., few unlinked loci may represent a relatively wide genomic region depending on the linkage relationships of the species. Obviously, the fewer the loci, the smaller the genomic region. Our results matches previous one for the Swedish model in (Sadedin et al., 2009) where ecotype formation was slower when the number of loci involved was larger and also recent results for the *L. saxatilis* Swedish system that shows the importance of adaptive recombination supression within inversions to facilitate ecotype evolution and adaptation (Koch et al., 2021). Moreover, recent work (Kautt et al., 2020) has highlighted that simple genetic architecture, as the one we modelled, not necessarily lead to speciation even with magic traits which is also consistent with our findings. It seems however that polygenic selection might be more efficient in driving sympatric speciation under some circumstances however this setting was out of the scope of the present study.

Most of the simulations evolved intermediate values of choosiness as a result of the trade-off between the effect of natural and sexual selection. Similar results have been obtained previously under some theoretical models and specific parameter range (see e.g. Rettelbach et al., 2013; Cotto and Servedio, 2017; Sachdeva and Barton, 2017 and references therein) and it is interesting that we have obtained the same result with a model for the specific case of *L. saxatilis* in the Galician coast using empirical estimates of the parameters (when available) and that the intermediate choosiness obtained in the simulations matches quite well empirical estimates of *L. saxatilis* and other littorinids (Fernández-Meirama et al., 2017a, 2017b; López-Cortegano et al., 2020).

Therefore, regarding our question number 2 on mating choice evolution as an effect of ecological speciation we may say that yes, under *L. saxatilis* Galician-like conditions, choosiness evolved during the process of ecological adaptation. Most of the evolved choosiness corresponded to positive assortative mating although negative assortative mating evolved under the combination of few loci and strong selection. As a general pattern, intermediate selection and few selective loci favoured the evolution of choosiness. However, this does not mean that the evolution of choosiness will irremediably cause ecological speciation. Actually, choosiness was typically maintained at intermediate values.

This was caused because there was an interplay between the evolution of assortative mating and local selective pressures. We modelled the evolution of choosiness as an ecologically neutral trait, although having as the target trait a magic trait, with realistic migration estimated for *L. saxatilis* in the Galician coast (Rolán-Alvarez et al., 2015a). These conditions imply that the evolution of choosiness is constrained by the maintenance of the trait polymorphism required for evolving local adaptation which is in turn affected by the current choosiness. If choosiness is strong, then rare, non optimal, phenotypes would mate between themselves. Therefore, during the process of local adaptation the alleles coding for strong choosiness are eliminated by viability selection. Under this setting, the highest amount of divergence occurs at intermediate choosiness values for a wide range of parameter values, which has been also observed in other studies (Rettelbach et al., 2013; Cotto and Servedio, 2017; Sachdeva and Barton, 2017). As far as we know, this is a new outcome regarding the evolution of mate choice for the *L. saxatilis* systems and is consistent with our intermediate estimates of assortative mating (Johannesson et al., 1995; Rolán-Alvarez et al., 1999, 2004; Cruz et al., 2004) and mate choice (Fernández-Meirama et al., 2017a, 2017b) for the Galician system. Positive assortative mating evolved in the three simulated habitats.

We also studied the correspondence between the preference based on the *c* trait and the assortative mating correlation measure. We already know that both measures matched well if the sampling is done at the correct scale where choice occurs (Estévez et al., 2018). However, we found important differences when comparing the choosiness values (recall that choosiness is a linear function of the *c* trait) with the value of the assortative mating correlation measure, especially when interpreted in the context of reinforcement. Reinforcement is defined in a broad sense as the evolution of traits that minimize hybrid formation in response to selection (Servedio et al., 2013; Pfennig, 2016). The effect of selection can be indirect i.e. reinforcement occurs due to selection against hybrids that indirectly results in prezygotic isolation or on the contrary, it involves the effect of natural or sexual selection acting directly on the prezygotic phenotypes involved in mate attraction. In the latter case, reinforcement is better called reproductive interference (Butlin and Ritchie, 2013; Hochkirch, 2013; Shaw and Mendelson, 2013).

Actually, we found higher mating correlation in the middle habitat than in the lower and upper ones. Therefore, if we use mating correlation as a proxy for assortative mating (Jiang et al., 2013; Janicke et al., 2019) we would say that we are detecting reproductive interference.

However, choosiness in the middle habitat, where hybrids can be formed, was very similar to choosiness in the other habitats, and so actually we cannot conclude that reproductive interference was evolved in our scenarios.

This putative reinforcement-like pattern cannot be due to a character displacement for the ecological trait (*x*) since there were no more extreme phenotypes in the middle area than in the other two. However, there was higher variation for the ecological trait in the middle habitat than in the others (coefficient of variation was at least one order of magnitude higher). Therefore, we should infer a higher sensitivity of the correlation measure to outliers (Pernet et al., 2013) which on the other hand would also explain the observed saturation effect of *r* over *c*. Actually, estimates from *L. saxatilis* empirical data indicate that mate choice has not increased in populations where the two ecotypes meet compared with those with only one ecotype (Fernández-Meirama et al., 2017a). All of the above corroborates that evolving high mate choice levels is difficult in presence of high gene flow. Therefore, regarding our question number 3 in the introduction, we may say that mate choice does not increase in the middle habitat under the simulated conditions of the Galician *L. saxatilis* system.

Finally, we considered the question number 4 about the impact of adding a cost to the mate choice and we observed in concordance with previous studies (Kopp and Hermisson, 2008), less reproductive isolation cases and just residual ecological speciation when the cost was added. As expected, when choosiness evolved its value was lower, but still relevant to have an ecological impact. In addition, there were an increasing importance on how the middle habitat was modelled and its impact on adaptation. When the middle habitat was neutral regarding the ecological trait, there were twice more adaptation failures than when the middle habitat was selective because when selection favours an intermediate phenotype this probably facilitated the transition to the optimal phenotypes for the lower shore (Schneider and Burger, 2006).

In summary, it seems that the *L. saxatilis* model system may correspond well to an incomplete ecological speciation case with intermediate choosiness and strong within habitat adaptation. The results obtained in our model system suggest that, the evolution of ecological speciation is not an immediate consequence of local divergent selection and mating preferences, but a fine tuning among the environmental conditions in the microhabitat and the hybrid middle zone, the genetic basis of the traits, the selection intensity and the mate choice stringency and cost. Similar results which have been obtained for other ecological speciation model cases (Safran et al., 2013; Cotto and Servedio, 2017) suggest that incomplete ecological speciation may be not rare.

These results may guide the search for new empirical data providing direction for future models to assist our understanding of the *L. saxatilis* system and ecological speciation in general. For example it would be interesting to check the particular condition that could facilitate higher values of choosiness and so putatively complete the ecological speciation process in this model system or even test whether the mate choice mechanism functions as a similarity-like mechanism as has been shown in other littorinids (Lau et al., 2021).

## Acknowledgements

We thank the reviewers for their constructive manuscript corrections. This work has received financial support from the Ministerio de Economía y Competitividad (BFU2013-44635-P; CGL2016-75482-P) and Xunta de Galicia (Grupo de Referencia Competitiva, ED431C 2020/05, Centro Singular de Investigación de Galicia accreditation 2019-2022), and by the European Union (European Regional Development Fund -ERDF). All authors declare to have no conflict of interest.

## References

Babik, W., Butlin, R. K., Baker, W. J., Papadopulos, A. S. T., Boulesteix, M., Anstett, M.-C., et al. (2009). How sympatric is speciation in the Howea palms of Lord Howe Island? Molecular Ecology 18, 3629–3638. doi:10.1111/j.1365-294X.2009.04306.x.

Barluenga, M., Stölting, K. N., Salzburger, W., Muschick, M., and Meyer, A. (2006). Sympatric speciation in Nicaraguan crater lake cichlid fish. Nature 439, 719. doi:10.1038/nature04325.

Barton, N. H. (2010). What role does natural selection play in speciation? Philosophical Transactions of the Royal Society B: Biological Sciences 365, 1825.

Berdahl, A., Torney, C. J., Schertzer, E., and Levin, S. A. (2015). On the evolutionary interplay between dispersal and local adaptation in heterogeneous environments. Evolution 69, 1390–1405. doi:10.1111/evo.12664.

Beverton, R. J. H., and Holt, S. J. (1957). On the dynamics of exploited fish populations. Ministry of Agriculture, Fisheries and Food (Series 2, Fyshery investigations 19.

Bolnick, D., and Fitzpatrick, B. M. (2007). Sympatric Speciation: Models and Empirical Evidence. Annual Review of Ecology, Evolution, and Systematics 38. Available at: citeulike-article-id:898435 http://dx.doi.org/10.1146/annurev.ecolsys.38.091206.095804.

Bolnick, D. I. (2004). Waiting for sympatric speciation. Evolution 58, 895–9.

Boulding, E. G. (1990). Are the opposing selection pressures on exposed and protected shores sufficient to maintain genetic differentiation between gastropod populations with high intermigration rates? Hydrobiologia 193, 41–52. doi:10.1007/BF00028065.

Boulding, E. G., Rivas, M. J., González Lavín, N., Rolán Alvarez, E., and Galindo, J. (2017). Size selection by a gape-limited predator of a marine snail: Insights into magic traits for speciation. Ecology and Evolution 7, 674–688. doi:10.1002/ece3.2659.

Butlin, R. K., and Ritchie, M. G. (2013). Pulling together or pulling apart: hybridization in theory and practice. Journal of Evolutionary Biology 26, 294–298. doi:10.1111/jeb.12080.

Butlin, R. K., Saura, M., Charrier, G., Jackson, B., Andre, C., Caballero, A., et al. (2014). Parallel Evolution of Local Adaptation and Reproductive Isolation in the Face of Gene Flow. Evolution 68, 935–949.

Carvajal-Rodríguez, A., and Rolán-Alvarez, E. (2014). A comparative study of Gaussian mating preference functions: a key element of sympatric speciation models. Biological Journal of the Linnean Society 113, 642–657.

Cotto, O., and Servedio, M. R. (2017). The Roles of Sexual and Viability Selection in the Evolution of Incomplete Reproductive Isolation: From Allopatry to Sympatry. The American Naturalist 190, 680–693. doi:10.1086/693855.

Coyne, J. A., and Orr, H. A. (2004). Speciation. Sinauer Available at: https://books.google.es/books?id=2Y9rQgAACAAJ.

Cruz, R., Carballo, M., Conde-Padin, P., and Rolan-Alvarez, E. (2004). Testing alternative models for sexual isolation in natural populations of Littorina saxatilis: indirect support for by-product ecological speciation? J Evol Biol 17, 288–93.

David, N., Fachada, N., and Rosa, A. C. (2017). “Verifying and Validating Simulations,” in Simulating Social Complexity: A Handbook Understanding Complex Systems., eds.B. Edmonds and R. Meyer (Cham: Springer International Publishing), 173–204. doi:10.1007/978-3-319-66948-9_9.

Debarre, F. (2012). Refining the conditions for sympatric ecological speciation. J Evol Biol 25, 2651–2660.

Debarre, F., and Gandon, S. (2011). Evolution in heterogeneous environments: between soft and hard selection. Am Nat 177, E84–97.

Dieckmann, U., and Doebeli, M. (1999). On the origin of species by sympatric speciation. Nature 400, 354–7.

Diz, A. P., Romero, M. R., Galindo, J., Saura, M., Skibinski, D. O. F., and Rolán-Alvarez, E. (2021). Proteomic analysis of F1 hybrids and intermediate variants in a Littorina saxatilis hybrid zone. Current Zoology. doi:10.1093/cz/zoab054.

Erlandsson, J., Rolan-Alvarez, E., and Johannesson, K. (1998). Migratory differences between ecotypes of the snail Littorina saxatilis on Galician rocky shores. Evolutionary Ecology 12, 913–924.

Estévez, D., Ng, T. P. T., Fernández-Meirama, M., Voois, J. M., Carvajal-Rodríguez, A., Williams, G. A., et al. (2018). A novel method to estimate the spatial scale of mate choice in the wild. Behavioral Ecology and Sociobiology 72, 195. doi:10.1007/s00265-018-2622-3.

Fernández-Meirama, M., Carvajal-Rodríguez, A., and Rolán-Alvarez, E. (2017a). Testing the role of mating preference in a case of incomplete ecological speciation with gene flow. Biological Journal of the Linnean Society 122, 549–557.

Fernández-Meirama, M., Estévez, D., Ng, T. P. T., Williams, G. A., Carvajal-Rodríguez, A., and Rolán-Alvarez, E. (2017b). A novel method for estimating the strength of positive mating preference by similarity in the wild. Ecology and Evolution 7, 2883–2893. doi:10.1002/ece3.2835.

Foote, A. D. (2018). Sympatric Speciation in the Genomic Era. Trends in Ecology & Evolution 33, 85–95. doi:10.1016/j.tree.2017.11.003.

Galindo, J., Martínez-Fernández, M., Rodríguez-Ramilo, S. T., and Rolán-Alvarez, E. (2013). The role of local ecology during hybridization at the initial stages of ecological speciation in a marine snail. J Evol Biol 26, 1472–1487. doi:10.1111/jeb.12152.

Gavrilets, S. (2004). Fitness landscapes and the origin of species. Princeton, N.J.: Princeton University Press.

Gavrilets, S., Vose, A., Barluenga, M., Salzburger, W., and Meyer, A. (2007). Case studies and mathematical models of ecological speciation. 1. Cichlids in a crater lake. Mol Ecol 16, 2893–909.

Getz, W. M., Salter, R., Seidel, D. P., and Hooft, P. (2016). Sympatric speciation in structureless environments. BMC Evolutionary Biology 16, 1–12. doi:10.1186/s12862-016-0617-0.

Hey, J. (2006). Recent advances in assessing gene flow between diverging populations and species. Current Opinion in Genetics & Development 16, 592–596.

Hochkirch, A. (2013). Hybridization and the origin of species. Journal of Evolutionary Biology 26, 247–251. doi:10.1111/j.1420-9101.2012.02623.x.

Janicke, T., Marie-Orleach, L., Aubier, T. G., Perrier, C., and Morrow, E. H. (2019). Assortative mating in animals and its role for speciation. The American Naturalist. doi:10.1086/705825.

Janson, K. (1987). Genetic drift in small and recently founded populations of the marine snail Littorina saxatilis. Heredity 58, 31.

Jiang, Y., Bolnick, D. I., and Kirkpatrick, M. (2013). Assortative Mating in Animals. The American Naturalist 181, E125–E138. doi:10.1086/670160.

Jiggins, C. D. (2006). Sympatric Speciation: Why the Controversy? Current Biology 16, R333–R334. doi:10.1016/j.cub.2006.03.077.

Johannesson, K., Panova, M., Kemppainen, P., Andre, C., Rolan-Alvarez, E., and Butlin, R. K. (2010). Repeated evolution of reproductive isolation in a marine snail: unveiling mechanisms of speciation. Philos Trans R Soc Lond B Biol Sci 365, 1735–47.

Johannesson, K., Rolán Alvarez, E., and Ekendahl, A. (1995). Incipient Reproductive Isolation Between Two Sympatric Morphs of the Intertidal Snail Littorina Saxatilis. Evolution 49, 1180–1190. doi:https://doi.org/10.1111/j.1558-5646.1995.tb04445.x.

Kautt, A. F., Kratochwil, C. F., Nater, A., Machado-Schiaffino, G., Olave, M., Henning, F., et al. (2020). Contrasting signatures of genomic divergence during sympatric speciation. Nature, 1–6. doi:10.1038/s41586-020-2845-0.

Kess, T., and Boulding, E. G. (2019). Genome-wide association analyses reveal polygenic genomic architecture underlying divergent shell morphology in Spanish Littorina saxatilis ecotypes. Ecol Evol 9, 9427–9441. doi:10.1002/ece3.5378.

Kess, T., Galindo, J., and Boulding, E. G. (2018). Genomic divergence between Spanish Littorina saxatilis ecotypes unravels limited admixture and extensive parallelism associated with population history. Ecology and Evolution 8, 8311–8327. doi:10.1002/ece3.4304.

Koch, E. L., Morales, H. E., Larsson, J., Westram, A. M., Faria, R., Lemmon, A. R., et al. (2021). Genetic variation for adaptive traits is associated with polymorphic inversions in Littorina saxatilis. Evolution Letters 5, 196–213. doi:10.1002/evl3.227.

Kopp, M., and Hermisson, J. (2008). Competitive speciation and costs of choosiness. Journal of Evolutionary Biology 21, 1005–1023. doi:10.1111/j.1420-9101.2008.01547.x.

Kopp, M., Servedio, M. R., Mendelson, T. C., Safran, R. J., Rodríguez, R. L., Hauber, M. E., et al. (2018). Mechanisms of Assortative Mating in Speciation with Gene Flow: Connecting Theory and Empirical Research. The American Naturalist 191, 1–20. doi:10.1086/694889.

Lau, S. L. Y., Williams, G. A., Carvajal-Rodríguez, A., and Rolán-Alvarez, E. (2021). An integrated approach to infer the mechanisms of mate choice for size. Animal Behaviour 175, 33–43. doi:10.1016/j.anbehav.2021.02.020.

Leinonen, T., McCairns, R. J. S., O’Hara, R. B., and Merilä, J. (2013). QST–FST comparisons: evolutionary and ecological insights from genomic heterogeneity. Nature Reviews Genetics 14, 179.

López-Cortegano, E., Carpena-Catoira, C., Carvajal-Rodríguez, A., and Rolán-Alvarez, E. (2020). Mate choice based on body size similarity in sexually dimorphic populations causes strong sexual selection. Animal Behaviour 160, 69–78. doi:10.1016/j.anbehav.2019.12.005.

Lotterhos, K. E., Moore, J. H., and Stapleton, A. E. (2018). Analysis validation has been neglected in the Age of Reproducibility. PLOS Biology 16, e3000070. doi:10.1371/journal.pbio.3000070.

Martin, C. H. (2013). Strong Assortative Mating By Diet, Color, Size, And Morphology But Limited Progress Toward Sympatric Speciation In A Classic Example: Cameroon Crater Lake Cichlids. Evolution 67, 2114–2123. doi:10.1111/evo.12090.

Martin, C. H., Cutler, J. S., Friel, J. P., Touokong, C. D., Coop, G., and Wainwright, P. C. (2015). Complex histories of repeated gene flow in Cameroon crater lake cichlids cast doubt on one of the clearest examples of sympatric speciation. Evolution 69, 1406–1422. doi:10.1111/evo.12674.

Massol, F., and Debarre, F. (2015). Evolution of dispersal in spatially and temporally variable environments: The importance of life cycles. Evolution 69, 1925–37.

Meyers, S. C., Liston, A., and Meinke, R. (2012). An Evaluation of Putative Sympatric Speciation within Limnanthes (Limnanthaceae). PLOS ONE 7, e36480. doi:10.1371/journal.pone.0036480.

Pérez-Figueroa, A., Cruz, F., Carvajal-Rodriguez, A., Rolan-Alvarez, E., and Caballero, A. (2005). The evolutionary forces maintaining a wild polymorphism of Littorina saxatilis: model selection by computer simulations. Journal of Evolutionary Biology 18, 191–202.

Perini, S., Rafajlović, M., Westram, A. M., Johannesson, K., and Butlin, R. K. (2020). Assortative mating, sexual selection, and their consequences for gene flow in Littorina. Evolution 74, 1482–1497. doi:10.1111/evo.14027.

Pernet, C. R., Wilcox, R., and Rousselet, G. A. (2013). Robust Correlation Analyses: False Positive and Power Validation Using a New Open Source Matlab Toolbox. Front Psychol 3, 606. doi:10.3389/fpsyg.2012.00606.

Pfennig, K. S. (2016). Reinforcement as an initiator of population divergence and speciation. Curr Zool 62, 145–154. doi:10.1093/cz/zow033.

Quesada, H., Posada, D., Caballero, A., Moran, P., and Rolan-Alvarez, E. (2007). Phylogenetic evidence for multiple sympatric ecological diversification in a marine snail. Evolution 61, 1600–12.

Reid, D. G. (1996). Systematics and evolution of Littorina. Available at: http://catalog.hathitrust.org/api/volumes/oclc/35218077.html.

Rettelbach, A., Kopp, M., Dieckmann, U., and Hermisson, J. (2013). Three Modes of Adaptive Speciation in Spatially Structured Populations. The American Naturalist 182, E215–E234.

Rettelbach, A., Servedio, M. R., and Hermisson, J. (2016). Speciation in peripheral populations: effects of drift load and mating systems. Journal of Evolutionary Biology 29, 1073–1090. doi:10.1111/jeb.12849.

Richards, E. J., Servedio, M. R., and Martin, C. H. (2019). Searching for Sympatric Speciation in the Genomic Era. BioEssays 41, 1900047. doi:10.1002/bies.201900047.

Richardson, J. L., Urban, M. C., Bolnick, D. I., and Skelly, D. K. (2014). Microgeographic adaptation and the spatial scale of evolution. Trends in Ecology & Evolution 29, 165–176.

Rolán-Alvarez, E. (2007). Sympatric speciation as a by-product of ecological adaptation in the Galician Littorina saxatilis hybrid zone. Journal of Molluscan Studies 73, 1–10. doi:10.1093/mollus/eyl023.

Rolán-Alvarez, E., Austin, C., and Boulding, E. G. (2015a). The contribution of the genus Littorina to the field of evolutionary ecology. Oceanography and Marine Biology: an Annual Review 53, 157–214.

Rolán Alvarez, E., Carballo, M., Galindo, J., Morán, P., Fernández, B., Caballero, A., et al. (2004). Nonallopatric and parallel origin of local reproductive barriers between two snail ecotypes. Molecular Ecology 13, 3415–3424. doi:https://doi.org/10.1111/j.1365-294X.2004.02330.x.

Rolán-Alvarez, E., Carvajal-Rodriguez, A., de Coo, A., Cortes, B., Estevez, D., Ferreira, M., et al. (2015b). The scale-of-choice effect and how estimates of assortative mating in the wild can be biased due to heterogeneous samples. Evolution 69, 1845–57.

Rolán Alvarez, E., Erlandsson, J., Johannesson, K., and Cruz, R. (1999). Mechanisms of incomplete prezygotic reproductive isolation in an intertidal snail: testing behavioural models in wild populations. Journal of Evolutionary Biology 12, 879–890.

Sachdeva, H., and Barton, N. H. (2017). Divergence and evolution of assortative mating in a polygenic trait model of speciation with gene flow. Evolution 71, 1478–1493. doi:10.1111/evo.13252.

Sadedin, S., Hollander, J., Panova, M., Johannesson, K., and Gavrilets, S. (2009). Case studies and mathematical models of ecological speciation. 3: Ecotype formation in a Swedish snail. Mol Ecol 18, 4006–23.

Safran, R. J., Scordato, E. S. C., Symes, L. B., Rodríguez, R. L., and Mendelson, T. C. (2013). Contributions of natural and sexual selection to the evolution of premating reproductive isolation: a research agenda. Trends in Ecology & Evolution 28, 643–650. doi:10.1016/j.tree.2013.08.004.

Savolainen, O., Lascoux, M., and Merila, J. (2013). Ecological genomics of local adaptation. Nat Rev Genet 14, 807.

Schneider, K. A., and Burger, R. (2006). Does competitive divergence occur if assortative mating is costly? J Evol Biol 19, 570–88.

Servedio, M. R., and Boughman, J. W. (2017). The Role of Sexual Selection in Local Adaptation and Speciation. Annual Review of Ecology, Evolution, and Systematics 48, 85–109. doi:10.1146/annurev-ecolsys-110316-022905.

Servedio, M. R., Hermisson, J., and Doorn, G. S. van (2013). Hybridization may rarely promote speciation. Journal of Evolutionary Biology 26, 282–285. doi:10.1111/jeb.12038.

Shaw, K. L., and Mendelson, T. C. (2013). The targets of selection during reinforcement. Journal of Evolutionary Biology 26, 286–287. doi:10.1111/jeb.12050.

Slatkin, M. (1995). A measure of population subdivision based on microsatellite allele frequencies. Genetics 139, 457–462. doi:10.1093/genetics/139.1.457.

Sokal, R. R., and Rohlf, F. J. (1981). Biometry. Second. New York: W. H. Freeman and Co.

Svardal, H., Rueffler, C., and Hermisson, J. (2015). A general condition for adaptive genetic polymorphism in temporally and spatially heterogeneous environments. Theoretical Population Biology 99, 76–97.

Thibert-Plante, X., and Gavrilets, S. (2013). Evolution of mate choice and the so-called magic traits in ecological speciation. Ecol Lett. Available at: http://www.ncbi.nlm.nih.gov/entrez/query.fcgi?cmd=Retrieve&db=PubMed&dopt=Citation&list_uids=23782866.

Thibert-Plante, X., and Hendry, A. P. (2009). Five questions on ecological speciation addressed with individual-based simulations. J Evol Biol 22, 109–23.

Thibert-Plante, X., and Hendry, A. P. (2011). The consequences of phenotypic plasticity for ecological speciation. Journal of Evolutionary Biology 24, 326–342.

Thiele, J. C., and Grimm, V. (2015). Replicating and breaking models: good for you and good for ecology. Oikos 124, 691–696.

Weissing, F., Edelaar, P., and van Doorn, G. S. (2011). Adaptive speciation theory: a conceptual review. Behavioral Ecology and Sociobiology 65, 461–480.

Westram, A. M., Rafajlović, M., Chaube, P., Faria, R., Larsson, T., Panova, M., et al. (2018). Clines on the seashore: The genomic architecture underlying rapid divergence in the face of gene flow. Evolution Letters 2, 297–309. doi:10.1002/evl3.74.

Wright, S. (1951). The genetical structure of populations. Annals Eugenics 15, 323–352.

