## Supplementary Tables for "A simulation study of the ecological speciation conditions in the Galician marine snail *Littorina saxatilis*"

* **Correspondence:**

Corresponding Author

**Table S1. Model parameter values for the comparison between Galician model and a symmetric model regarding carrying capacity and migration.**

| | | | Value | |
|---|---|---|---|---|
| | **Parameter** | **Symbol** | **Galician** | **Symmetric** |
| Demography | Generation number | *T* | 20,000 | 20,000 |
| | **Number of exposed demes** | | **4** | **12** |
| | Number of intermediate demes | | 2 | 2 |
| | **Number of sheltered demes** | | **20** | **12** |
| | **Per deme carrying capacity in exposed habitat** | *K* | **15,000** | **5481** |
| | **Per deme carrying capacity in intermediate and sheltered** | | **3750** | **5481** |
| | Mean offspring number | *b* | 50 | 50 |
| | Between deme migration probability | *m* | | |
| | | 0 demes | 0.75 | 0.75 |
| | | 1 demes | 0.15 | 0.15 |
| | | 2 demes | 0.1 | 0.1 |
| Genome Structure | Number of microsatellites | | *L* = 8 | *L* = 8 |
| | Ecological magic trait (*L* loci) | *x* | *L* = 4 | *L* = 4 |
| | Male mating trait (*L* loci) | *c* | *L* = 4 | *L* = 4 |
| Selection | Selection strength | $\sigma_s$ | 0.45 | 0.45 |
| | Habitat optimum selection in middle | θ | no (neutral) | no (neutral) |
| Mating | Evaluated females per male | $N_f$ | 10 | 10 |
| | Mating preference tolerance | $\sigma_a$ | 0.05 | 0.05 |
| | Mate choice cost | | no | no |
| Mutation | Neutral mutation rate per locus | $\mu_0$ | $10^{-3}$ | $10^{-3}$ |
| | Trait mutation rate per locus | μ | $10^{-5}$ | $10^{-5}$ |

**Table S2. Results for the ecological trait and choosiness under Galician and Symmetric models for the parameters given in Table S1 i.e. *L*= 4 loci, intermediate selection strength $\sigma_s$ = 0.45, and tolerance $\sigma_a$ = 0.05 and neutral model in the middle habitat. Values are averages of 100 replicates plus minus the standard error of the mean.**

| Model | x trait value in lower shore | x trait value in middle shore | Choosiness (*C*) | | |
|---|---|---|---|---|---|
| | | | **Upper** | **Middle** | **Lower** |
| Galician | 0.0005± 0.0002 | 0.15 ± 0.004 | 0.52±0.019 | 0.54±0.016 | 0.54 ±0.018 |
| Symmetric | 0.007 ± 0.0001 | 0.48 ± 0.007 | 0.58±0.018 | 0.58±0.015 | 0.56±0.015 |

**Table S3. Cases with evolution of negative assortative mating defined as *C*<-0.1 (c<0.45). Values are computed over 20 replicates. Empty cells imply no colonization of the lower shore.**

| $\sigma_s$ | $\sigma_a$ | Middle zone | *L* | % runs with negative *C* | | | Mean *C* | | |
|---|---|---|---|---|---|---|---|---|---|
| | | | | **Lower** | **Middle** | **Upper** | **Lower** | **Middle** | **Upper** |
| 0.15 | 0.1 | selective | 4 | 65% | 75% | 80% | -0.32 | -0.36 | -0.38 |
| 0.15 | 0.1 | neutral | 4 | 50% | 50% | 55% | -0.21 | -0.07 | -0.12 |
| 0.15 | 0.05 | selective | 4 | 90% | 95% | 95% | -0.5 | -0.55 | -0.51 |
| 0.15 | 0.05 | neutral | 4 | 80% | 80% | 95% | -0.27 | -0.26 | -0.42 |
| 0.15 | 0.1 | selective | 8 | 15% | 25% | 35% | 0.08 | 0.08 | -0.02 |
| 0.15 | 0.1 | neutral | 8 | --- | 25% | 20% | --- | 0.04 | 0.05 |
| 0.15 | 0.05 | selective | 8 | 0% | 0% | 5% | 0.1 | 0.18 | 0.1 |
| 0.15 | 0.05 | neutral | 8 | --- | 5% | 0% | --- | 0.14 | 0.14 |

**Table S4. Results for the Galician model for the parameters given in Table S1 i.e. *L*= 4 loci, intermediate selection strength $\sigma_s$ = 0.45, and tolerance $\sigma_a$ = 0.05 but comparing the choosiness in the two middle habitat scenarios i.e. neutral versus selective. Values are averages of 100 replicates plus minus the standard error of the mean.**

| Middle Scenario | Choosiness (*C*) | | |
|---|---|---|---|
| | **Upper** | **Middle** | **Lower** |
| Neutral | 0.52±0.02 | 0.54±0.02 | 0.54 ±0.02 |
| Selective (θ=0.5) | 0.52±0.02 | 0.53±0.02 | 0.53±0.02 |

**Table S5. Results for the Galician model for the same parameters as in Table S4 but comparing the mating correlation instead of choosiness. Values are averages of 100 replicates plus minus the standard error of the mean.**

| Middle Scenario | Correlation (*r*) | | |
|---|---|---|---|
| | Upper | Middle | Lower |
| Neutral | 0.63±0.04 | 0.85±0.01 | 0.79 ±0.04 |
| Selective (θ=0.5) | 0.63±0.04 | 0.87±0.02 | 0.69±0.04 |

**Table S6. Results for mating correlation under the Galician and the Symmetric model for the parameters given in Table S1 i.e. *L*= 4 loci, intermediate selection strength $\sigma_s$ = 0.45, and tolerance $\sigma_a$ = 0.05 and neutral model in the middle habitat. Values are averages of 100 replicates plus minus the standard error of the mean.**

| Model | Correlation (*r*) | | |
|---|---|---|---|
| | Upper | Middle | Lower |
| Galician | 0.63±0.04 | 0.85±0.01 | 0.79 ±0.04 |
| Symmetric | 0.73±0.04 | 0.99±0.002 | 0.81±0.04 |

**Table S7. Percentage of speciation, defined as mate choice trait c ≥ 0.9 in the middle habitat and $Q_{ST}$ ≥ 0.9, for the scenarios with high ($\sigma_s$ =0.15), intermediate ($\sigma_s$ =0.45) or low selection ($\sigma_s$ =1) under neutral or selective middle habitat. Tolerance was $\sigma_a$ = 0.05. *L* is the number of loci for the ecological and male mating traits. Values are averages of 100 replicates.**

| $\sigma_s$ | Middle zone | *L* | % speciation |
|---|---|---|---|
| 0.15 | neutral | 4 | 0% |
| 0.15 | neutral | 16 | 0% |
| 0.15 | selective | 4 | 0% |
| 0.15 | selective | 16 | 0% |
| 0.45 | neutral | 4 | 11% |
| 0.45 | neutral | 16 | 0% |

| | | | |
|---|---|---|---|
| 0.45 | selective | 4 | 11% |
| 0.45 | selective | 16 | 0% |
| 1 | neutral | 4 | 15% |
| 1 | neutral | 16 | 0% |
| 1 | selective | 4 | 13% |
| 1 | selective | 16 | 0% |

**Table S8. ANOVA results for the scenarios with mating cost, showing % of variance explained by each factor for the ecological (*x*) and choice (*c*) traits, mating correlation (*r*), population size (*N*) and differentiation measures ($F_{ST}$ and $Q_{ST}$). Only the two most important factor interactions are presented. The values within cells correspond to the % at lower, middle, upper shore, by this order, except for differentiation. The asterisk indicates significance at the 0.001 level.**

| **Factor** | **Response** | | | | | |
|---|---|---|---|---|---|---|
| | ***x*** | ***c*** | ***r*** | ***N*** | $F_{ST}$ | $Q_{ST}$ |
| $\sigma_s$ | 67*, 27*, 74* | 39, 45, 47* | 12, 85*, 87* | 29, 20*, 23 | 65* | 14* |
| $\sigma_a$ | 0, 0, 3 | 0, 1, 0 | 11, 2, 0 | 6, 0, 0 | 3 | 0 |
| **L** | 11*, 11*, 2 | 10, 3, 0 | 1, 4, 0 | 7, 7*, 7 | 6* | 14* |
| $\theta$ | 0, 8*, 3 | 24, 6, 2 | 17, 1, 0 | 1, 23*, 1 | 1 | 14* |
| L× $\sigma_s$ | 10, 14*, 2 | 4, 8, 12 | 8, 1, 4 | 1, 10*, 7 | 4* | 14* |
| $\theta \times \sigma_s$ | 0, 15*, 5 | 1, 10, 8 | 2, 0, 1 | 0, 19*, 7 | 7* | 14* |
